# Loss of dihydroceramide desaturase drives neurodegeneration by disrupting endoplasmic reticulum and lipid droplet homeostasis in glial cells

**DOI:** 10.1101/2024.01.01.573836

**Authors:** Yuqing Zhu, Kevin Cho, Haluk Lacin, Yi Zhu, Jose T. DiPaola, Beth A. Wilson, Gary J. Patti, James B. Skeath

**Affiliations:** Department of Genetics, Washington University School of Medicine, 4523 Clayton Avenue, St. Louis, MO 63110, USA; Department of Chemistry, Washington University in St. Louis, One Brookings Drive, St. Louis, MO 63130, USA; Department of Medicine, Washington University School of Medicine, St. Louis, MO 63110, USA; Center for Mass Spectrometry and Metabolic Tracing, Washington University in St. Louis, One Brookings Drive, St. Louis, MO 63130, USA; Division of Biological and Biomedical Systems, University of Missouri-Kansas City, Kansas City, MO 64110, USA

**Keywords:** *DEGS1*, ceramide, sphingolipids, neurodegeneration, glia, lipid droplet, endoplasmic reticulum, leukodystrophy

## Abstract

Dihydroceramide desaturases convert dihydroceramides to ceramides, the precursors of all complex sphingolipids. Reduction of DEGS1 dihydroceramide desaturase function causes pediatric neurodegenerative disorder hypomyelinating leukodystrophy-18 (HLD-18). We discovered that *infertile crescent (ifc)*, the *Drosophila DEGS1* homolog, is expressed primarily in glial cells to promote CNS development by guarding against neurodegeneration. Loss of *ifc* causes massive dihydroceramide accumulation and severe morphological defects in cortex glia, including endoplasmic reticulum (ER) expansion, failure of neuronal ensheathment, and lipid droplet depletion. RNAi knockdown of the upstream ceramide synthase *schlank* in glia of *ifc* mutants rescues ER expansion, suggesting dihydroceramide accumulation in the ER drives this phenotype. RNAi knockdown of *ifc* in glia but not neurons drives neuronal cell death, suggesting that *ifc* function in glia promotes neuronal survival. Our work identifies glia as the primary site of disease progression in HLD-18 and may inform on juvenile forms of ALS, which also feature elevated dihydroceramide levels.

## INTRODUCTION

Sphingolipids are key structural and functional components of the cell membrane in all eukaryotic cells and are enriched in glial cells, such as oligodendrocytes, where they comprise up to 30% of all membrane lipids.^1^ All complex sphingolipids, like glycosylsphingolipids and sphingomyelin, derive from ceramide.^2–4^ In the *de novo* ceramide biosynthesis pathway, dihydroceramide desaturases, such as DEGS1, produce ceramide from dihydroceramide by catalyzing the formation of a *trans* double bond between carbons 4 and 5 of the sphingoid backbone, which enhances conformational plasticity.^5,6^ In humans, bi-allelic mutations in *DEGS1* cause hypomyelinating leukodystrophy-18 (HLD-18), a progressive, often fatal pediatric neurodegenerative disease marked by cerebral atrophy, white matter reduction, and hypomyelination.^7–9^ The primary neural cell type impacted by loss of *DEGS1* function and the cell biology of how disruption of ceramide synthesis leads to neurodegeneration remain unknown.

The *de novo* ceramide biosynthetic pathway is well conserved among higher metazoans.^4,10–13^ *De novo* ceramide synthesis occurs in the endoplasmic reticulum (ER) and starts with the rate-limiting activity of the serine palmitoyltransferase (SPT) complex, which condenses serine and palmitoyl-CoA (lauoryl-CoA in flies) to form 3-Ketosphinganine, which is converted to sphinganine by 3-Ketosphinganine reductase. Ceramide synthases condense sphinganine with acyl-CoA to generate dihydroceramide, which is converted to ceramide by dihydroceramide desaturases. Ceramide is then efficiently transported by the specific ceramide transporter CERT from the ER to the Golgi,^14^ where it undergoes headgroup modifications to produce complex sphingolipids that eventually translocate to the plasma membrane.^15^ Mutations in most members of the SPT complex, ceramide synthases, and DEGS1 lead to neurodegeneration,^4,12,13^ identifying the *de novo* ceramide biosynthesis pathway as a hotspot for neurodegenerative disease mutations.

Consistent with its central role in ceramide biogenesis, reduction or loss of *DEGS1* function in human patients or cell lines, mice, zebrafish, and flies drive dihydroceramide accumulation and ceramide depletion.^8,9,16,17^ HLD-18 patients display reduced myelin sheath thickness in peripheral nerves, and knockdown of *DEGS1* function in zebrafish reduces the number of myelin basic protein-positive oligodendrocytes,^8,9^ suggesting *DEGS1* regulates Schwann cell and oligodendrocyte development. In *Drosophila*, genetic ablation of *infertile crescent* (*ifc*), the fly *DEGS1* ortholog, drives activity-dependent photoreceptor degeneration,^17^ suggesting that *ifc* function is crucial for neuronal homeostasis. In addition, forced expression of a wildtype *ifc* transgene in neurons, glia, or muscles was shown to rescue the *ifc* mutant phenotype.^17^ Whether *ifc/DEGS1* acts primarily in glia, neurons, or other cells to regulate nervous system development then remains to be determined.

Research on *DEGS1* points to dihydroceramide accumulation as the driver of nervous system defects caused by *DEGS1* deficiency. Pharmacological or genetic inhibition of ceramide synthase function, which should reduce dihydroceramide levels, suppresses the observed reduction of MBP-positive oligodendrocytes in zebrafish and the activity-dependent photoreceptor degeneration in flies triggered by reduction of *DEGS1/ifc* function.^9,17^ Dihydroceramide accumulation may also contribute to other neurodegenerative diseases, as gain of function mutations in components of the SPT complex cause juvenile amyotrophic lateral sclerosis due to elevated sphingolipid biosynthesis, with dihydroceramides showing the greatest relative increase of all sphingolipids.^18–22^ How dihydroceramide accumulation alters the cell biology of neurons, glia, or both to trigger myelination defects and neurodegeneration is unclear.

With its vast genetic toolkit, *Drosophila* is a powerful system in which to dissect the genes and pathways that regulate glial development and function.^23,24^ In flies, six morphologically and functionally distinct glial subtypes regulate nervous system development and homeostasis.^25,26^ The surface perineurial and subperineurial glia act as physiochemical barriers to protect the nervous system and control metabolite exchange with the hemolymph, carrying out a similar function as the human blood-brain-barrier.^27,28^ Residing between the surface glia and the neuropil, cortex glia ensheathe the cell bodies of neuroblasts and neurons in the CNS in protective, nutritive, honeycomb-like membrane sheaths. Ensheathing glia define the boundary of the neuropil and insulate axons, dendrites, and synapses from neuronal cell bodies in the CNS.^29^ Astrocyte-like glia extend fine membrane protrusions that infiltrate the neuropil and form a meshwork of cellular processes that ensheathe synapses and regulate synaptic homeostasis in the CNS.^29^ In the PNS, wrapping glia reside internally to the surface glia and insulate axons in the PNS to enhance neuronal signaling and provide energy support.

Using the *Drosophila* model, we found that *ifc* acts primarily in glia to regulate CNS development with its loss disrupting glial morphology. Our work supports a model in which inappropriate accumulation and retention of dihydroceramide in the ER drives ER expansion, glial swelling, and the failure of glia to enwrap neurons, ultimately leading to neuronal degeneration as a secondary consequence of glial dysfunction. Given the conserved nature of *de novo* ceramide biosynthesis, our findings likely illuminate the exact mechanism through which elevated dihydroceramide levels drive neuronal degeneration and cell death in flies and humans.

## RESULTS

### *ifc* contributes to the regulation developmental timing and CNS structure

In an EMS-based genetic screen, we uncovered three non-complementing mutations that when homozygous or trans-heterozygous to each other resulted in identical phenotypes, including a 3-day or greater delay in reaching the late-third larval instar stage, reduced brain size, progressive ventral nerve cord elongation, axonal swelling, and lethality at the late larval or early pupal stage (Figure 1A; data not shown). Whole genome sequencing revealed that *ifc* was independently mutated in each line: *ifc^js^*^1^ and *ifc^js2^* encode V276D and G257S missense mutations, respectively, and *ifc^js3^* encodes a W162* nonsense mutation (Figures 1A and 1B). Sanger sequencing also uncovered a missense mutation, E241K, in the molecularly uncharacterized *ifc^1^* allele (Figure 1B). All four mutations reside in the fatty acid desaturase domain, the hotspot for mutations in human *DEGS1* that cause HLD-18 (Figures 1B-B’).

**Figure 1.**
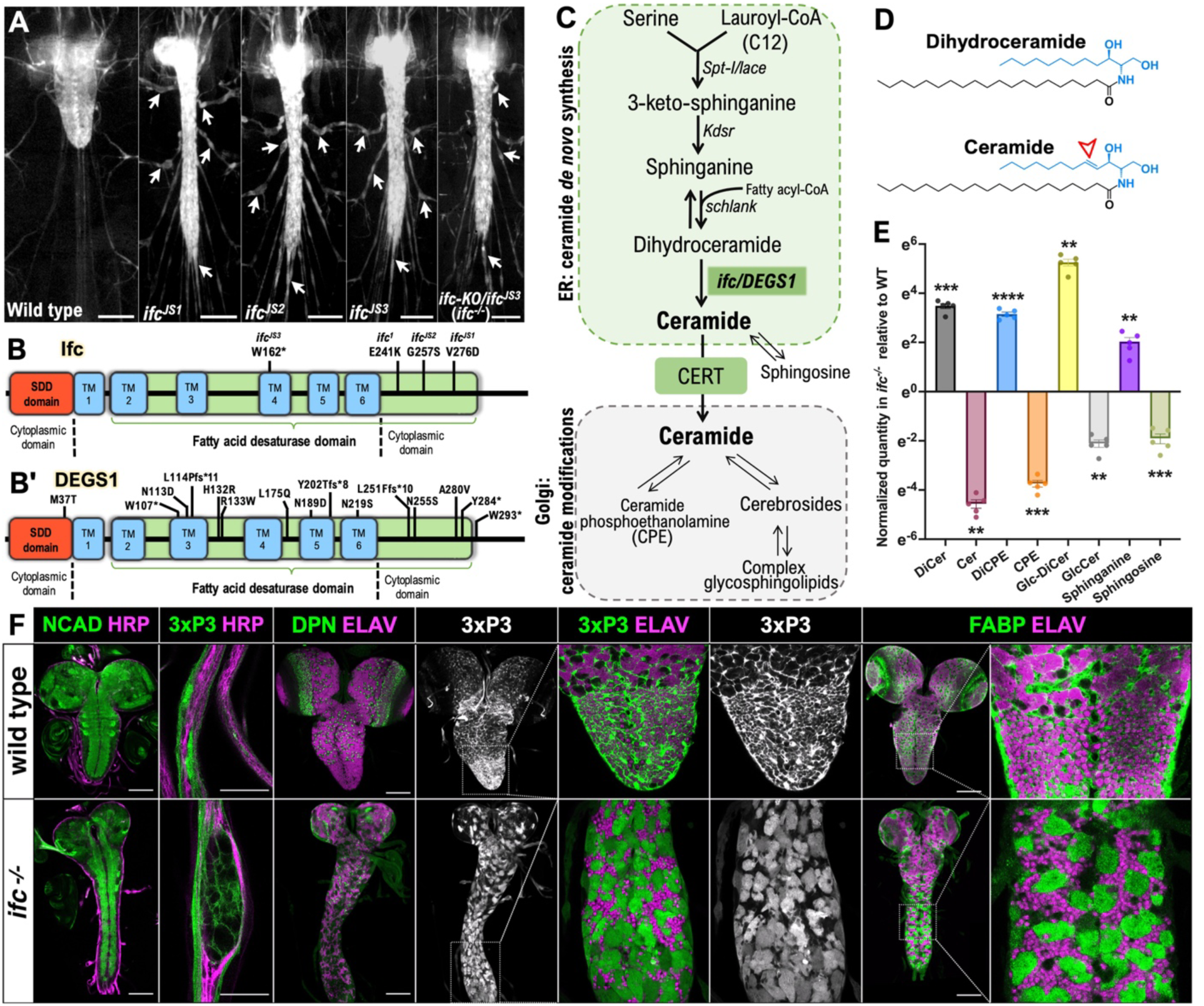
*ifc* regulates CNS and glial morphology. A) Ventral views of late-third instar larvae of indicated genotype showing 3xP3 RFP labeling of CNS and nerves. Arrowheads indicate nerve bulges; scale bar is 200μm. B-B’) Schematic of Ifc (B) and human DEGS1 (B’) proteins indicating location and nature of *ifc* mutations and 15 HLD-18-causing *DEGS1* mutations.^7,8,9^ C) Schematic of *de novo* ceramide biosynthesis pathway indicating the subcellular location of ceramide synthesis and ceramide modifications. D) Chemical structure of dihydroceramide and ceramide; arrow indicates *trans* carbon-carbon double bond between C4 and C5 in the sphingoid backbone created by the enzymatic action of Ifc/DEGS1. E) Normalized quantification of the relative levels of dihydroceramide, ceramide, and six related sphingolipid species in the dissected CNS of wild-type and *ifc^-/-^* late-third instar larvae. F) Ventral views of *Drosophila* CNS and peripheral nerves in wild-type and *ifc^-/-^* mutant late-third instar larvae labeled for NCAD to mark the neuropil, HRP to label axons, RFP to label glia, Dpn to label neuroblasts, ELAV to label neurons, and FABP to label cortex glia. Anterior is up; scale bar is 100μm for whole CNS images and 20μm for peripheral nerve image. Statistics: * denotes *p* < 0.05, ** denotes *p* < 0.01, *** denotes *p* < 0.001, **** *denotes p* < 0.0001.

As a prior study reported that a CRISPR-generated, gene-specific deletion of *ifc, ifc-KO,* resulted in early larval lethality^17^, we first confirmed the genetic nature of our *ifc* alleles because they caused late larval-early pupal lethality. Complementation crosses of each *ifc* allele against a deficiency of the region (Df(2L)BSC184) and *ifc-KO* revealed that all combinations, including larvae trans-heterozygous for *ifc-KO* over Df(2L)BSC184, survived to the late larval-early pupal stage and yielded phenotypes identical to those detailed above for the newly uncovered *ifc* alleles. Flies homozygous for *ifc-KO*, however, died as early larvae. Further analysis uncovered second site mutation(s) in the 21E2 chromosomal region responsible for the early lethal phenotype of the *ifc-KO* chromosome (see Methods). When uncoupled from these mutation(s), larvae homozygous for the “clean” *ifc-KO* chromosome developed to the late larval-early pupal stage and manifested phenotypes identical to the other *ifc* alleles. This analysis defined the correct lethal phase for *ifc* and identified our *ifc* alleles as strong loss of function mutations.

### Loss of *ifc* function drives ceramide depletion and dihydroceramide accumulation

As *ifc/DEGS1* converts dihydroceramide to ceramide, we used untargeted lipidomics on whole larvae and the isolated CNS from wild-type and *ifc-KO/ifc^JS3^* larvae (hereafter termed *ifc^-/-^* larvae) to assess the effect of loss of *ifc* function on metabolites in the ceramide pathway (Figure 1C). Loss of *ifc* function resulted in a near complete loss of ceramides and a commensurate increase in dihydroceramides in the CNS and whole larvae (Figures 1E, S1). Sphinganine, the metabolite directly upstream of dihydroceramide, also exhibited a significant increase in its levels in the absence of *ifc* function, while metabolites further upstream were unchanged in abundance or undetectable (Figures 1E, S1). Ceramide derivatives like sphingosine, CPE, and Glucosyl-Ceramide (Glc-Cer), were reduced in levels and replaced by their cognate dihydroceramide forms (e.g., Glc-DiCer) (Figures 1E, S1). Loss of the enzymatic function of Ifc then drives dihydroceramide accumulation and ceramide loss.

### *ifc* governs glial morphology and survival

To connect this metabolic profile to a cellular phenotype, we assayed *ifc* function in the CNS. We leveraged the expression of fatty acid binding protein (FABP) as a marker of cortex glia^30^ and that of the *M{3xP3-RFP.attP}* phi-C31 “landing pad” transgene, which resided in the isogenic target chromosome of our screen, as a marker of most glia (Figures S2 and S3). In addition, we labeled neuroblasts with Deadpan (Dpn), neurons with ELAV, and axons with N-Cadherin (NCad). In *ifc*^-/-^ larvae, we observed a clear reduction in Dpn-positive neuroblasts in the optic lobe, swelling of glia in peripheral nerves, enhanced RFP expression in the CNS, and the presence of large swollen, cortex glia identified by RFP labeling and fatty acid binding protein (FABP) expression (Figures 1F, S4, and S5). In wild-type larvae, cortex glia display compact cell bodies and fully enwrap individual neuronal cell bodies with their membrane sheaths (Figures 1F and S4).^30^ In *ifc*^-/-^ larvae, cortex glia display swollen cell bodies, fail to fully enwrap neuronal cell bodies, displace neurons from their regular arrangement, and appear to contain brightly fluorescent RFP-positive aggregates (Figures 1F, S6). *ifc* is then necessary for glial development and function in the larval nervous system. We observed identical CNS phenotypes in larvae homozygous mutant for the *ifc^js1^*and *ifc^js2^* alleles (Figure S7).

To track the impact of *ifc* on glial morphology, we combined GAL4 lines specific for each glial subtype with a UAS-linked membrane-tagged GFP transgene (Myr-GFP) and the MultiColor FlpOut system.^31–33^ Using this approach, we determined that loss of *ifc* function affects all CNS glial subtypes except perineurial glia (Figures 2E-E’ and 2J-J’). Cortex glia appeared swollen, failed to enwrap neurons, and accumulated large amounts of Myr-GFP^+^ internal membranes (Figures 2A-A’ and 2F-F’). Ensheathing glia (Figures 2B-B’ and 2G-G’) and subperineurial glia (Figures 2D-D’ and 2I-I’) also displayed swollen, disorganized cell bodies and accumulated Myr-GFP^+^ internal membranes. Astrocyte-like glia displayed smaller cell bodies, reduced membrane extensions, and disrupted organization along the dorsal ventral nerve cord (Figures 2C-C’). We conclude that *ifc* regulates the morphology of most glial subtypes in the larval CNS.

**Figure 2.**
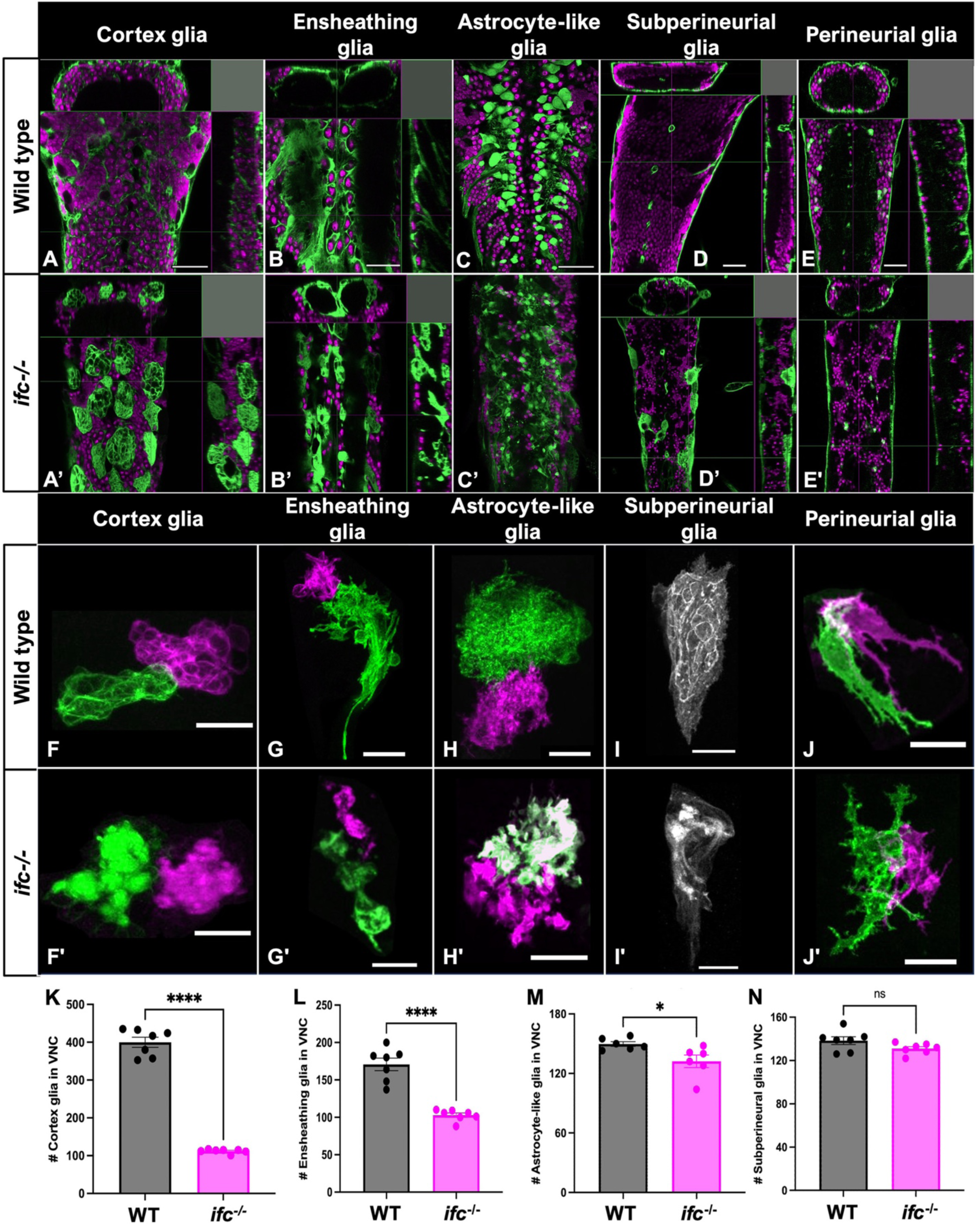
Loss of *ifc* disrupts glial morphology. A-E’) High magnification ventral views and X-Z and Y-Z projections of the nerve cord of wildtype and *ifc^-/-^* late third instar larvae labeled for ELAV (magenta) for neurons and Myr-GFP (green) for cell membranes of indicated glial subtype. Anterior is up; scale bar is 40μm. F-J’) High magnification views of individual glial cells of indicated glial subtype in the nerve cord of wildtype and *ifc^-/-^* larvae created by the MultiColor-FlpOut method.^33^ Anterior is up; scale bar is 20μm. K-N) Quantification of total number of indicated glial subtype in the nerve cord of wildtype and *ifc^-/-^* late third instar larvae (n = 7 for K, L, N; n = 6 for M). Statistics: * denotes *p* < 0.05, **** denotes *p* < 0.0001, and ns, not significant.

Next, we asked if loss of *ifc* function alters the number of each glial subtype. Using the same glial subtype-specific GAL4 lines to drive a nuclear-localized GFP transgene, we counted the total number of all CNS glial subtypes, except perineurial glia, in wild type and *ifc^-/-^* larvae. To remove the small size of the brain in *ifc^-/-^* larvae as a confounding factor, we focused our analysis on the ventral nerve cord. The number of subperineurial glia was unchanged between the two genotypes, but we observed a 12%, 40%, and 72% reduction in the number of astrocyte-like, ensheathing, and cortex glia, respectively, in *ifc*^-/-^ larvae relative to wild-type (Figures 2K-N). Our data reveal a broad role for *ifc* in regulating glial cell morphology and number in the *Drosophila* larval CNS. Subsequent experiments revealed that a reduction in cell proliferation and an increase in apoptosis both contribute to the observed reduction in the number of cortex glia (Figure S8).

### *ifc* acts in glia to regulate glial and CNS development

Prior work in zebrafish showed that *DEGS1* knockdown reduced the number of myelin basic protein-positive oligodendrocytes;^9^ in flies, loss of *ifc* function in the eye drove photoreceptor degeneration.^17^ Neither study uncovered the cell type in which *ifc/DEGS1* acts to regulate neural development. To address this question, we used the GAL4/UAS system, RNAi-mediated gene depletion, and gene rescue approaches to see if *ifc* acts in neurons or glia to control glial development and CNS morphology. First, we used a UAS-linked *ifc*-RNAi transgene to deplete *ifc* function in all neurons (*elav-GAL4*; *repo-GAL80*) or all glia (*repo-GAL4*). Focusing on FABP-positive cortex glia due to their easily scorable phenotype, we found that pan-glial, but not pan-neuronal, knockdown of *ifc* recapitulated the swollen cortex glia phenotype observed in *ifc* mutant larvae (Figures 3C-D and 3J).

**Figure 3.**
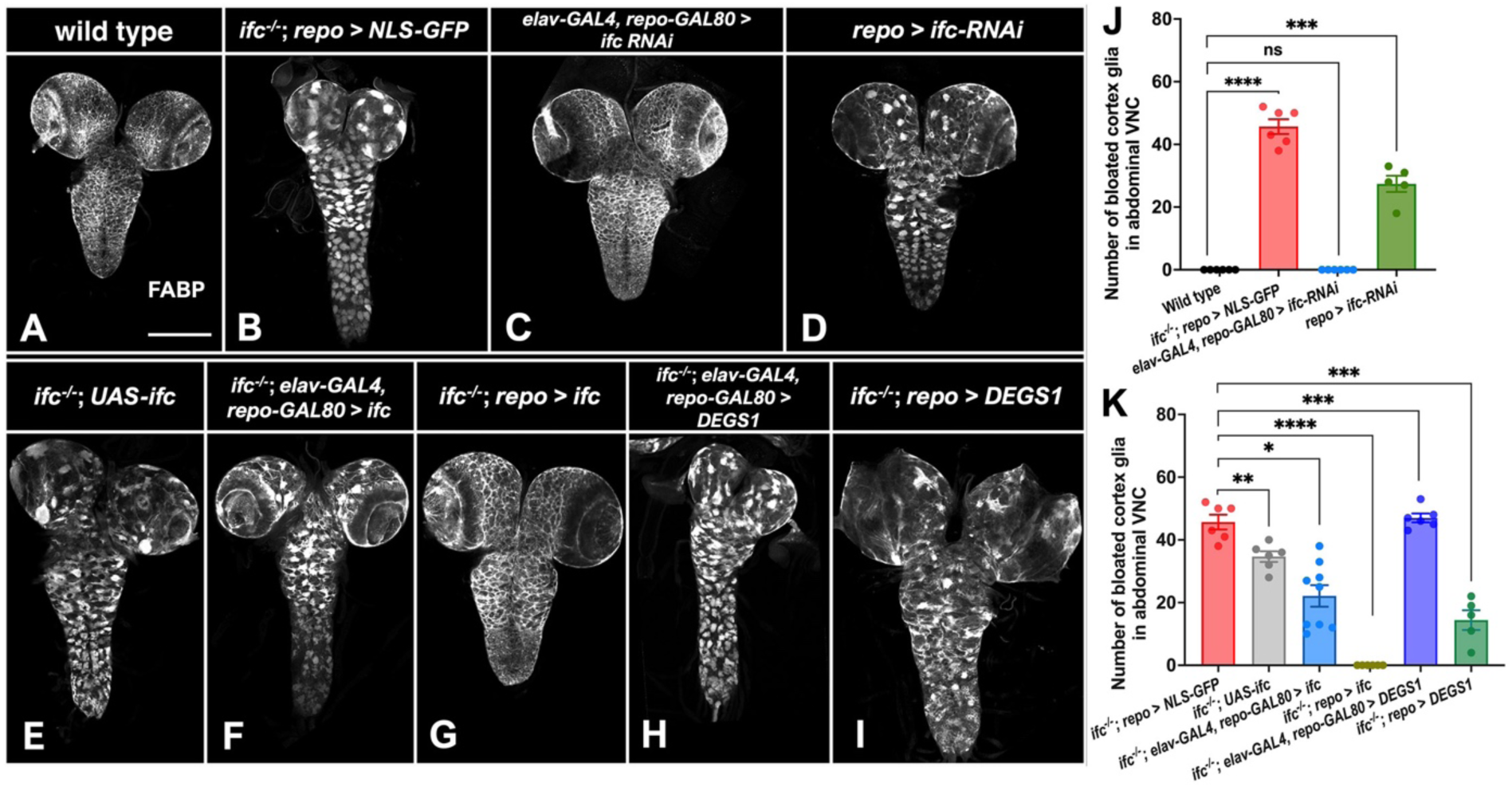
*ifc* acts in glia to regulate CNS structure and glial morphology. A-I) Ventral views of photomontages of the CNS of late third instar larvae labeled for FABP (greyscale) to mark cortex glia in late third instar larvae of indicated genotype. Neuronal-specific transgene expression was achieved by using *elav-GAL4* combined with *repo-GAL80*; glial-specific transgene expression was achieved by using *repo-GAL4*. J-K) Quantification of the number of swollen cortex glia in the abdominal segments of the CNS of late-third instar larvae of the indicated genotype for the RNAi (J) and gene rescue assays (K). Statistics: * denotes *p* < 0.05, ** denotes *p* < 0.01, *** denotes *p* < 0.001, **** denotes *p* < 0.0001, and ns, not significant.

To complement our RNAi approach, we asked if GAL4-driven expression of a wild-type *Drosophila ifc* or human *DEGS1* transgene rescued the *ifc^-/-^* CNS phenotype. In the absence of a GAL4 driver, the *ifc* transgene drove weak rescue of the cortex glia phenotype (Figures 3E and 3K), consistent with modest GAL4-independent transgene expression reported for UAS-linked transgenes.^34^ Pan-neuronal expression of *ifc* drove modest rescue of the *ifc* CNS phenotype beyond that observed for the *ifc* transgene alone (Figures 3F and 3K), but pan-glial expression of *ifc* fully rescued the *ifc* mutant cortex glia phenotype and other CNS phenotypes (Figures 3H, 3K, and **S9**). Identical experiments using the human *DEGS1* transgene revealed that only pan-glial *DEGS1* expression provided rescuing activity, albeit at much weaker levels than the *Drosophila ifc* transgene (Figures 3G, 3I, and 3K). Pan-glial expression of the *Drosophila ifc* transgene was, in fact, ∼15-fold more potent than the human *DEGS1* transgene at rescuing the *ifc* lethal phenotype to adulthood: When *ifc* was expressed in all glia, 57.9% of otherwise *ifc* mutant flies survived to adulthood (n=2452), but when *ifc* was replaced by *DEGS1* only 3.9% of otherwise *ifc* mutant flies reached adulthood (n=1303). No *ifc* mutant larvae reached adulthood in the absence of either transgene (n=1030). We infer that *ifc* acts primarily in glia to govern CNS development and that human *DEGS1* can partially substitute for *ifc* function in flies despite a difference in the preferred length of the sphingoid backbone in flies versus mammals.^10^

Results of our gene rescue experiments conflict with a prior study on *ifc* in which expression of *ifc* in neurons was found to rescue the *ifc* phenotype.^17^ In this context, we note that *elav-GAL4* drives UAS-linked transgene expression not just in neurons, but also in glia at appreciable levels,^35, 36^ and thus needs to be paired with *repo-GAL80* to restrict GAL4-mediated gene expression to neurons. Thus, “off-target” expression in glial cells may account for the discrepant results. It is, however, more difficult to reconcile how neuronal or glial expression of *ifc* would rescue the observed lethality of the *ifc-KO* chromosome given the presence additional lethal mutations in the 21E2 region of the second chromosome.

### *ifc* is predominately expressed in glia and localizes to the ER

Next, we tracked *ifc* expression in the CNS via RNA in-situ hybridization and an *ifc-T2A-GAL4* transcriptional reporter. RNA in-situ hybridization revealed that *ifc* is widely expressed in the CNS (Figure 4A), most obviously in the distinctive star-shaped astrocyte-like glia (Figure 4B), which are marked by Ebony expression.^37^ RNA in-situ hybridization was not ideal for tracing *ifc* expression in other cells, likely due to signal diffusion. Thus, we paired the *ifc*-*T2A-GAL4* transcriptional reporter with a nuclear RFP (nRFP) transgene^38^ and confirmed authenticity of the *ifc-T2A-GAL4* line by its strong expression in astrocyte-like glia (Figure 4C). Using this approach, we observed strong nRFP expression in all glial cells (Figures 4D and S10A) and modest nRFP expression in all neurons (Figures 4E and S10B), suggesting *ifc* is transcribed at higher levels in glial cells than neurons in the larval CNS.

**Figure 4.**
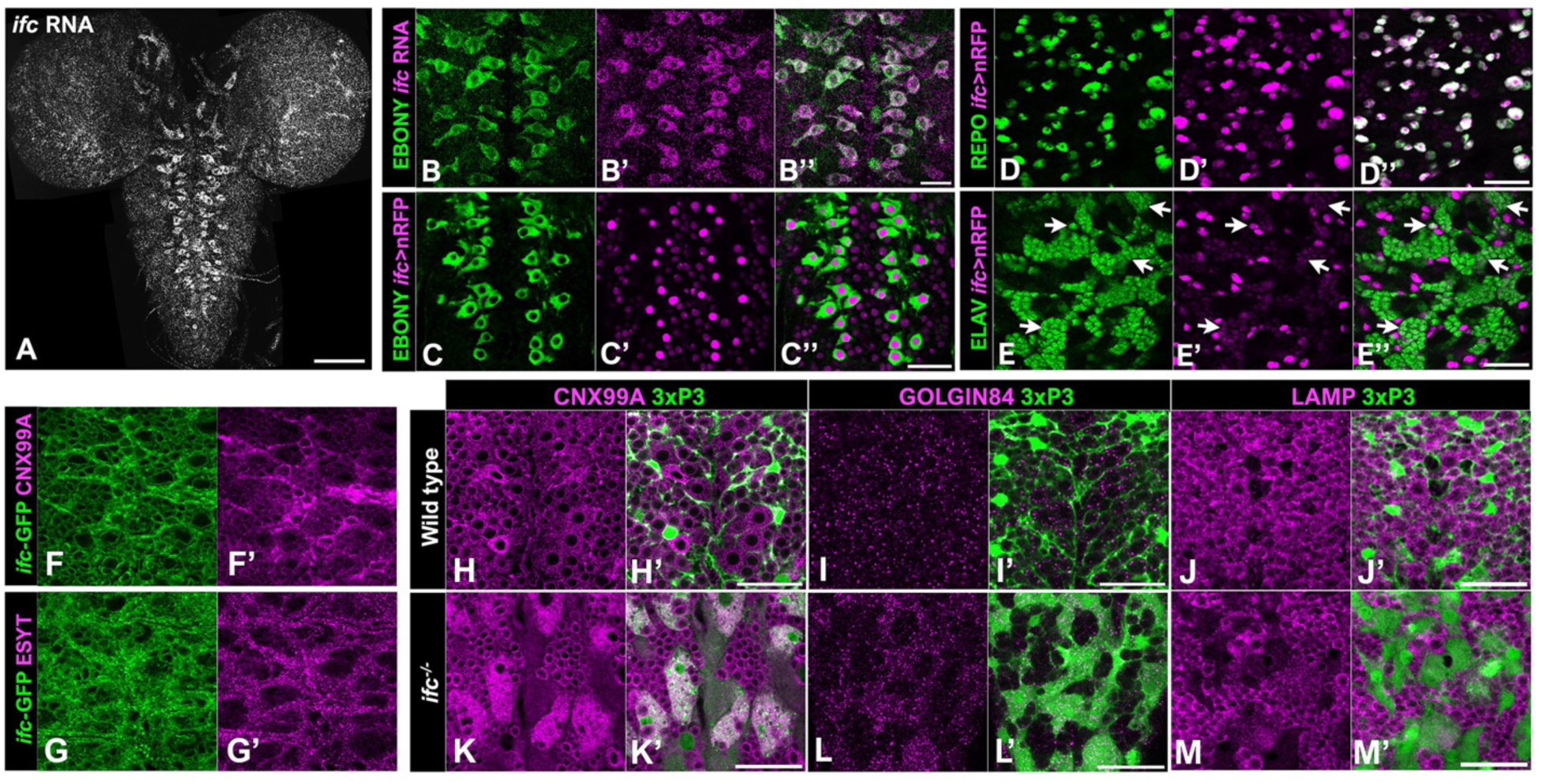
Loss of ifc drives ER expansion in cortex glia. A-E) Dorsal (A, B-B’’, and C-C’’) and ventral (D-D’’ and E-E’’) views of the CNS of late-third instar wild-type larvae labeled for *ifc* RNA (grey in A; magenta in B’), *ifc-GAL4>nRFP* (magenta; C’-E’), EBONY to mark astrocytes (green; B and C), REPO to mark glia (green; D), and ELAV to mark neurons (green; E). Panels D-D’’ and E-E’’ show surface and interior views, respectively, along the Z-axis on the ventral side of the nerve cord. Arrowheads in E-E’’ identify neurons with low level *ifc-GAL4* expression. F-G) High magnification ventral views of thoracic segments in the CNS of wild-type late third instar larvae labeled for GFP (green; F and G), CNX99A (magenta; F’), ESYT (magenta; G’). H-M) Late third instar larvae of indicated genotype labeled for 3xP3-RFP (green; H’-M’), CNX99A (magenta; H and K), GOLGIN84 (magenta; I and L), and LAMP (magenta; J and M). Anterior is up; scale bar is 100μm for panel A and 30μm for panels B-M.

Using a fosmid transgene that harbors a GFP-tagged version of *ifc* flanked by ∼36-kb of its endogenous genomic region and molecular markers for ER, cis-Golgi, and trans-Golgi,^39–42^ we found that the Ifc-GFP colocalized strongly with the ER markers Calnexin 99A (CNX99A) and ESYT (Figures 4F-F’, 4G-G’, and S7) and weakly with the cis-Golgi marker GOLGIN84 (Figure S10C) and the trans-Golgi marker GOLGIN245 (Figure S10D). Our results indicate that Ifc localizes primarily to the ER, aligning with the presumed site of *de novo* ceramide biosynthesis and prior work on *DEGS1* localization in cell lines.^8,43^

### Loss of *ifc* drives ER expansion and lipid droplet loss in cortex glia

We next asked if loss of *ifc* function altered ER, Golgi, or lysosome morphology. Focusing on cortex glia, we observed a clear expansion of the ER marker CNX99A (Figures 4H-H’ and K-K’), a mild enrichment of the Golgi markers, GOLGIN84 and GOLGIN245, in diffuse “clouds” (Figures 4I-I’, 4L-L’, S10E-F), and a reduction in expression of the lysosome marker LAMP (Figures 4J-J’ and M-M’) in *ifc^-/-^* larvae. The expansion of ER markers in *ifc* mutant larvae compelled us to obtain high-resolution views of organelle structure in cortex glia via transmission electron microscopy (TEM). In wildtype, cortex glia display a compact cytoplasm that surrounds a large nucleus (Figures 5A-B) and extend glial sheaths that fully enwrap adjacent neuronal cell bodies (black arrows; Figures 5C-D). In *ifc^-/-^* larvae, cortex glia display enlarged cell bodies with a maze-like pattern of internal membranes (solid white arrows; Figures 5A’-B’ and 5E-E’) and fail to enwrap neurons (hollow white arrows; Figures 5C’-D’). The internal membrane structures appear to assume an ER-like identity, as we observed significant overlap between the ER marker CNX99A and the membrane marker Myr-GFP when Myr-GFP was driven by a cortex glia-specific GAL4 line (Figure 5F). We observed similar yet milder effects on cell swelling and internal membrane accumulation in subperineurial and wrapping glia in abdominal nerves (purple and pink shading, respectively; Figures 5G-G’ and 5H-H’), indicating that loss of *ifc* drives internal membrane accumulation in and swelling of multiple glial subtypes.

**Figure 5.**
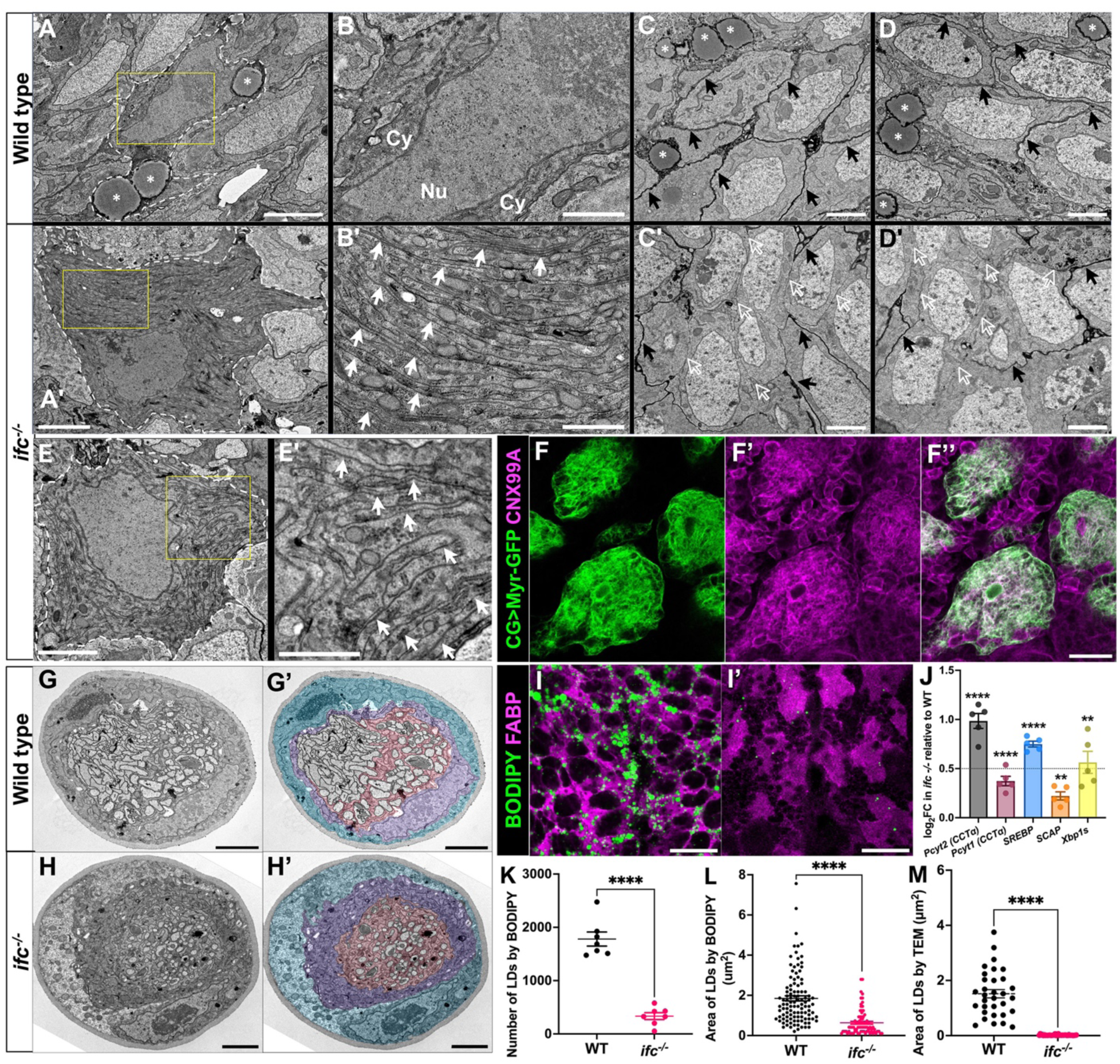
Loss of *ifc* leads to internal membrane accumulation and lipid droplet loss in cortex glia. A-E’) TEM images of cortex glia cell body (A-A’ and B-B’) and neuronal cell bodies (C-C’ and D-D’) at low (A-A’) and high (B-B’, C-C’, and D-D’) magnification in the nerve cord of wildtype (A-D) and *ifc^-/-^* (A’-D’ and E-E’) late third instar larvae. (A-A’) Dotted lines demarcate cell boundary of cortex glia; yellow squares highlight regions magnified in B, B’, and E’. Scale bar is 3μm for A and A’ and 1μm for B-B’. (B-B’) Cy denotes cytoplasm; Nu denotes nucleus. Solid white arrows highlight the layered internal membranes that occupy the cytoplasm of *ifc^-/-^* cortex glia. (C-C’ and D-D’) Black arrows highlight cortex glia membrane extensions that enwrap neuronal cell bodies; hollow white arrows denote the absence of cortex glia membrane extensions; white asterisk denotes lipid droplets. Scale bar is 2μm. E-E’) An additional example of membrane-filled cortex glia cell body in *ifc^-/-^* larvae. Scale bar is 2μm for E and 1μm for E’. F) Cortex glia in *ifc* mutant larvae labeled for Myr-GFP (green) to label membranes and CNX99A to label ER membranes. Scale bar is 30μm. G-H) Black and white and colored TEM cross-sections of peripheral nerves in wild type and *ifc^-/-^* late-third instar larvae. Blue marks perineurial glia; purple marks subperineurial glia; pink marks wrapping glia. Scale bar: 2μm. I-I’) High magnification ventral views of abdominal segments in the ventral nerve cord of wild-type (I) and *ifc* mutant (I’) third instar larvae labeled for BODIPY (green) to mark lipid droplets and FABP (magenta) to label cortex glia. Anterior is up; scale bar is 30μm. J) Graph of log-fold change of transcription of five genes that promote membrane lipid synthesis in *ifc^-/-^* larvae relative to wildtype. Dotted line indicates a log2 fold change of 0.5 in the treatment group compared to the control group. -M) Quantification of the number (G) and area of lipid droplets (H and I) in the dissected CNS of wildtype and *ifc^-/-^* larvae. Statistics: * denotes *p* < 0.05, ** denotes *p* < 0.01, *** denotes *p* < 0.001, **** denotes *p* < 0.0001, and ns, not significant.

TEM analysis also revealed a near complete depletion of lipid droplets in the CNS of *ifc*^-/-^ larvae (compare Figures 5A, 5C, and 5D to 5A’, 5C’ and 5D’; lipid droplets marked by asterisk; Figure 5M), which we confirmed using BODIPY to mark neutral lipids in FABP-positive cortex glia (Figures 5I-L). In the CNS, lipid droplets form primarily in cortex glia^30^ and are thought to contribute to membrane lipid synthesis through their catabolism into free fatty acids versus acting as an energy source in the brain.^44^ Consistent with the possibility that increased membrane lipid synthesis drives lipid droplet reduction, RNA-seq assays of dissected nerve cords revealed that loss of *ifc* drove transcriptional upregulation of genes that promote membrane lipid biogenesis, such as *SREBP*, the conserved master regulator of lipid biosynthesis, *SCAP*, an activator of SREBP, and *Pcyt1/Pcyt2,* which promote phosphatidylcholine and phosphatidylethanolamine synthesis.^45–47^ The spliced form of *Xbp-1* mRNA (*Xbp-1s*) (Figure 5J), which promotes membrane lipid synthesis required for ER biogenesis and is activated by the unfolded protein response (UPR),^48^ is also upregulated in the CNS of *ifc* mutant larvae. Most ER chaperones, which are typically transcriptionally upregulated upon UPR activation,^49^ were however downregulated (Figure S11A), suggesting that in this case misfolded protein is not the factor that triggers UPR activation and increased ER membrane biogenesis upon loss of *ifc*.

### Loss of *ifc* increases the saturation levels of triacylglycerols and membrane phospholipids

Lipid droplets are composed largely of triacylglycerols (TGs),^47^ and we observed a 5-fold and 3-fold drop in TG levels in the CNS and whole larvae of *ifc* mutant larvae relative to wild-type (Figure 6A). Our lipidomics analysis also revealed a shift of TGs toward higher saturation levels in the absence of *ifc* function (Figures 6A-C). Consistent with this, we observed transcriptional upregulation of most genes in the Lands cycle, which remodels phospholipids by replacing existing fatty acyl groups with new fatty acyl groups (Figures S11B-C).^50–53^ As TG breakdown results in free fatty acids that can be used for membrane phospholipid synthesis, we asked if changes in TG levels and saturation were reflected in the levels or saturation of the membrane phospholipids phosphatidylcholine (PC), phosphatidylethanolamine (PE), and phosphatidylserine (PS). In the absence of *ifc* function, PC and PE exhibited little change in quantity (Figures 6D, 6G), but the levels of the less abundant PS were increased three-fold increase in the CNS of *ifc* mutant larvae relative to wild-type (compare Figure 6J to Figures 6D and 6G). All three phospholipids, however, displayed increased saturation levels: The relative levels of all PC, PE, and PS species were reduced, except for the most saturated form of each phospholipid – 18:1/18:1 – which is increased (Figures 6E-F and 6H-L). This increase was more pronounced in the CNS than whole larvae (Figures 6F, 6I, and 6L), implying that loss of *ifc* function creates greater demand for lipid remodeling in the CNS.

**Figure 6.**
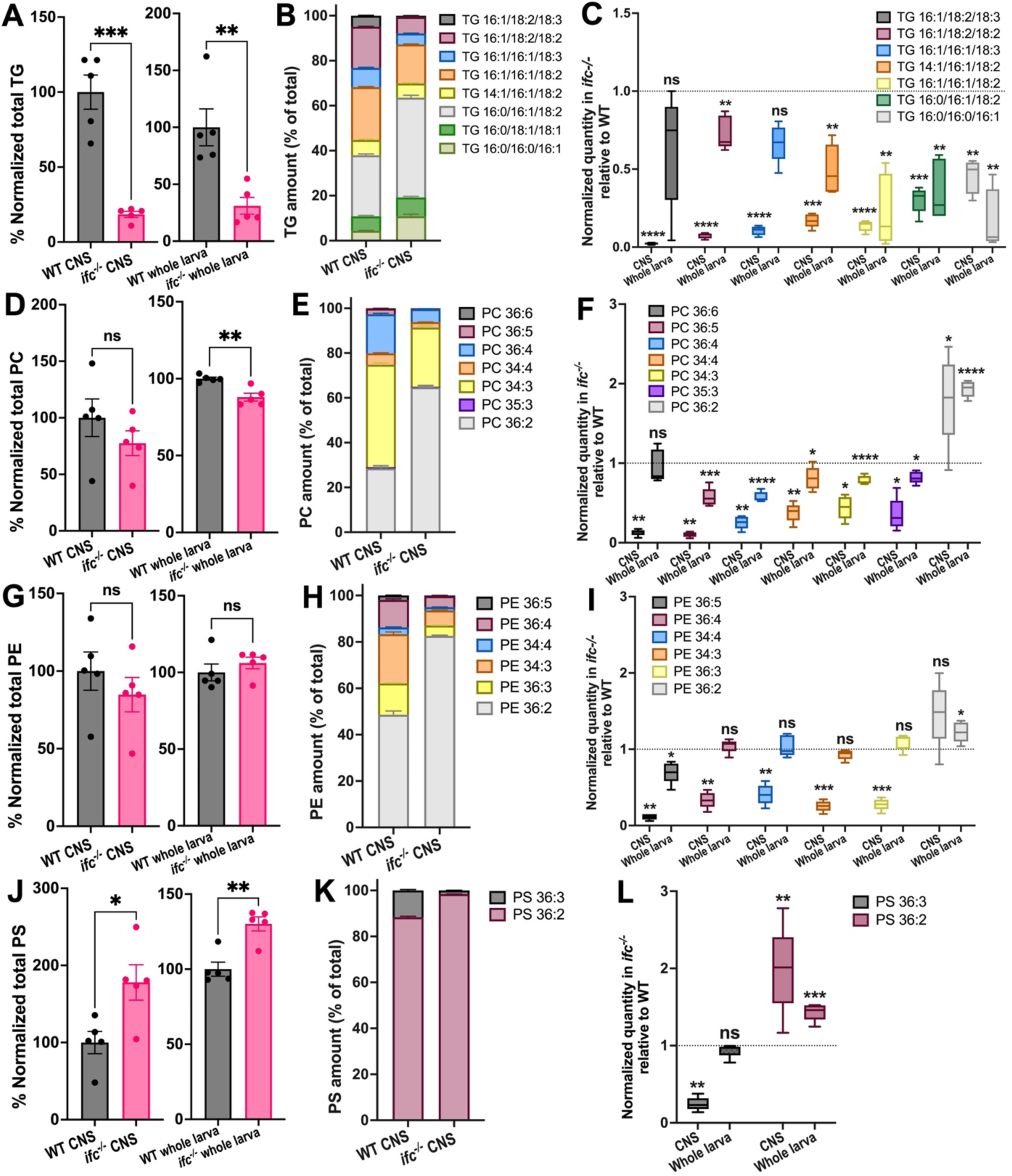
PC, PE, PS, and TG exhibit higher saturation levels in a CNS-specific manner in *ifc* mutant late third instar larvae. A-C) Quantification of total (A) and species-specific (B and C) TGs in whole larvae (A and C) and dissected CNS (A, B, and C) of wildtype and *ifc^-/-^*larvae. D-F). Quantification of total (D) and species-specific (E and F) PCs in whole larvae (D and F) and dissected CNS (D, E, and F) of wildtype and *ifc^-/-^*larvae. G-I) Quantification of total (G) and species-specific (H and I) PEs in whole larvae (G and H) and dissected CNS (G, H, and I) of wild-type and *ifc^-/-^* larvae. J-L) Quantification of total (J) and species-specific (K and L) PSs in the whole larvae (J and L) and dissected CNS (J, K, and L) of wild-type and *ifc^-/-^* larvae. Statistics: * denotes *p* < 0.05, ** denotes *p* < 0.01, *** denotes *p* < 0.001, **** denotes *p* < 0.0001, and ns, not significant.

### Reduction of dihydroceramide synthesis suppresses the *ifc* CNS phenotype

Following its synthesis, ceramide is transported by CERT from the ER to the Golgi, but CERT is less efficient at transporting dihydroceramide than ceramide.^13^ The expanded nature of the ER in *ifc* mutant larvae supports a model in which loss of *ifc* triggers excessive accumulation and retention of dihydroceramide in the ER due to CERT’s inefficiency of transporting dihydroceramide to the Golgi, driving ER expansion and glial swelling. If this model is correct, reduction of dihydroceramide levels should suppress the *ifc* CNS phenotype. In agreement with this model, we found that the *schlank^G0365^* loss of function allele dominantly suppressed the enhanced RFP expression (Figure 7M) and CNS elongation phenotypes of *ifc* (Figure 7N). We also found that glial-specific depletion of *schlank* suppressed the internal membrane accumulation (Figures 7A-B), reduced lipid droplet size (Figures 7C-D, and 7O), glial swelling (Figures 7E-F, and S12), enhanced RFP expression (Figure 7M), CNS elongation (Figure 7N), and reduced optic lobe (Figure S5) phenotypes observed in otherwise *ifc* mutant larvae. Our data support the model that inappropriate retention of dihydroceramide in the ER drives ER expansion and glial swelling and dysfunction.

**Figure 7.**
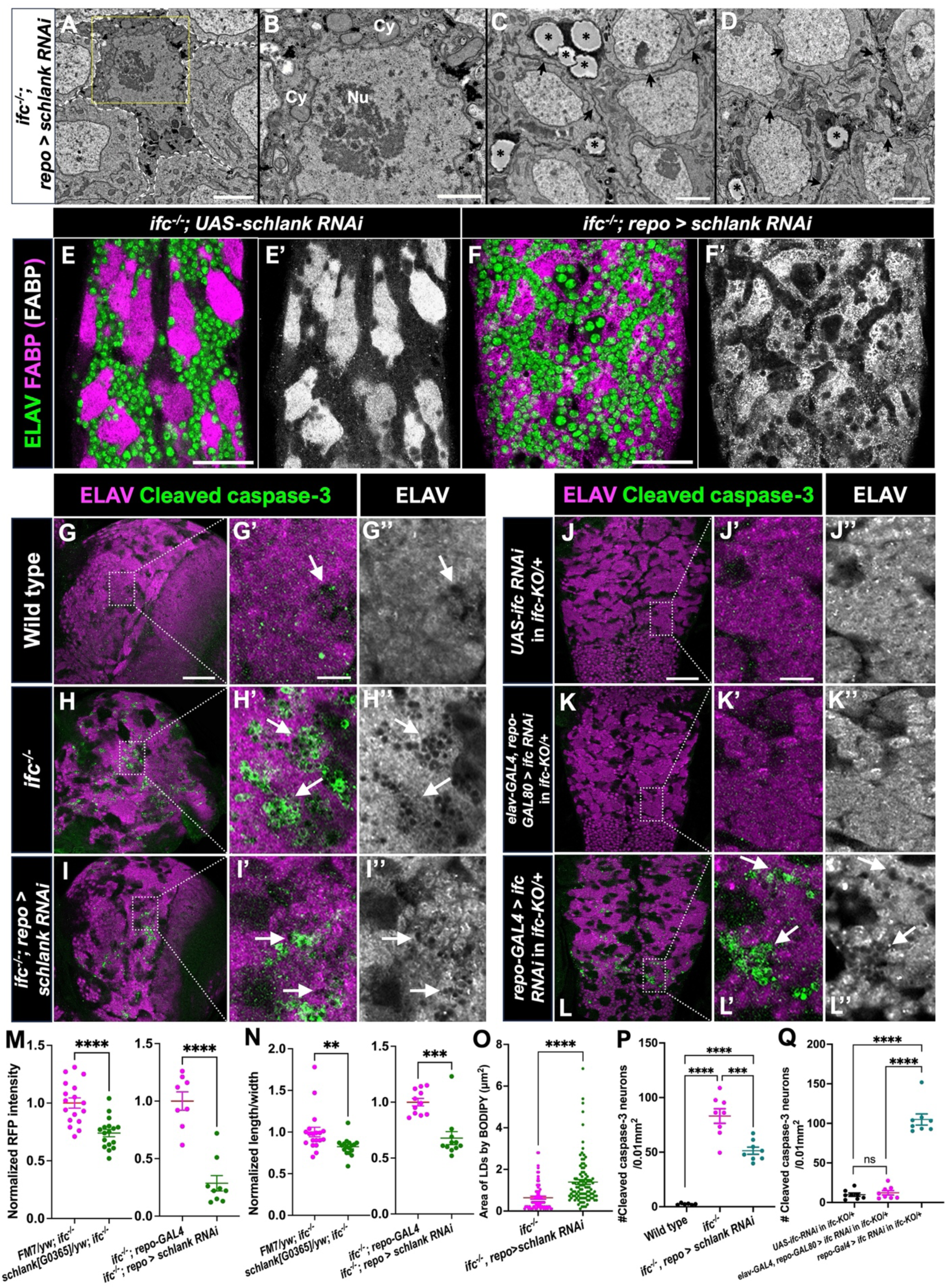
Glial-specific knockdown of *ifc* triggers neuronal cell death. A-D) TEM images of cortex glia cell body (A and B) and neuronal cell bodies (C and D) at low (A) and high (B, C, and D) magnifications in the nerve cord of *ifc^-/-^*; *repo>schlank RNAi* late third instar larvae. Dotted lines demarcate cell boundary of cortex glia; yellow squares highlight regions magnified in A. Scale bar: 3μm for A, 1μm for B, and 2μm for C and D. (B) Cy denotes cytoplasm; Nu denotes nucleus. (C-D) Black asterisk denotes lipid droplets. E-F) Ventral views of abdominal sections of CNS of *ifc^-/-^*; *UAS-schlank RNAi/+* larvae (E) and *ifc^-/-^*; repoGAL4/*UAS-schlank RNAi* larvae (F) labeled for neurons (ELAV, Green) and cortex glia (FABP, magenta/grey). Scale bar is 30μm for E-F. G-L) Low (G-L) and high (G’-L’ and G’’-L’’) magnification views of the brain (G-I) and nerve cord (J-L) of late-third instar larvae of the indicated genotypes labeled for ELAV (magenta or greyscale) and Caspase-3 (green). Arrows indicate regions of high Caspase-3 signal and/or apparent neuronal cell death identified by perforations in the neuronal cell layer. Scale bar is 50μm for panels G-L and 10μm for panels G’-L’’. M-N) Quantification of CNS elongation (M) and 3xP3 RFP intensity (N) in *ifc* mutants alone, *ifc* mutants with one copy of *schlank[G0365]* loss-of-function allele, or *ifc* mutants in which *schlank* function is reduced via RNAi in glial cells. O) Quantification of the area of lipid droplets in dissected CNS of *ifc* mutants and *ifc* mutants in which *schlank* function is reduced via RNAi in glial cells. Anterior is up in all panels. P-Q) Quantification of Cleaved Caspase-3 neurons for panels G-I (P) and J-L (Q). Statistics: * denotes *p* < 0.05, ** denotes *p* < 0.01, *** denotes *p* < 0.001, **** denotes *p* < 0.0001, and ns, not significant.

### Glial-specific depletion of *ifc* function triggers neuronal cell death

Loss of *DEGS1/ifc* in human and flies has been shown to promote neurodegeneration and neuronal cell death,^8,9,17^ but whether neuronal death results from an intrinsic defect in neurons or is induced by glial dysfunction remains unclear. By using Cleaved caspase-3 as a marker of dying cells^54,55^ and ELAV to track neurons, we tracked neuronal cell death in the brain and ventral nerve cord of wild-type and *ifc^-/-^*mutant late-third instar larvae. In wildtype, little to no cell death was evident in the brain or nerve cord and neurons appear smoothly packed next to each other; in contrast, in *ifc^-/-^* larvae, significant cell death was apparent in the brain and to a lesser degree in the nerve cord, with Caspase-3 staining often highlighting small perforations in the neuronal cell layer (Figures 7G-H, 7P, and S7). This perforated pattern was associated with and more expansive than Caspase-3 staining, indicating the perforations mark Caspase-positive dying neurons and Caspase-negative dead neurons. Glial-specific depletion of *schlank* function in otherwise *ifc^-/-^* larvae suppressed the neuronal cell death phenotype (Figures 7I and 7P), supporting the model that loss of *ifc* function specifically in glia, rather than a specific requirement for *ifc* function in neurons, drives neuronal cell death. To directly test whether *ifc* function in glia is required to guard against neuronal cell death, we used the GAL4-UAS system and RNAi mediated gene interference to deplete *ifc* function specifically in glia (*repo-GAL4*) or in neurons (*elav-GAL4; repo-GAL80*) and found that glial-specific, but not neuronal-specific, depletion of *ifc* function drove significant neuronal cell death in the brain and to a greater extent the nerve cord, a phenotype that was enhanced upon removal of one copy of *ifc* (Figures 7J-L, 7Q, and S13). Loss of *ifc* function then triggers glial dysfunction, which in turn drives neuronal cell death.

## DISCUSSION

Our work on *ifc* supports a model in which loss of *ifc/DEGS1* function drives glial dysfunction through the accumulation and inappropriate retention of dihydroceramide in the ER of glia with this proximal defect driving ER expansion, glial swelling, and subsequent neuronal cell death and neurodegeneration. A large increase in dihydroceramide would also impact ER membrane structure: ER membranes are typically loosely packed, thin, semi-fluid structures, with sphingolipids comprising just 3% of ER phospholipids.^47^ Sphingolipids in general and dihydroceramide in specific are highly saturated lipids, promote lipid order, tighter lipid packing, membrane rigidity, and thicker membranes.^56,57^ Our observation of thick ER membranes in cortex glia in *ifc*^-/-^ larvae (Figure 5) aligns with increased dihydroceramide levels in the ER. In this context, we note that of many clinical observations made on an HLD-18 patient, one was widening of the ER in Schwann cells,^8^ a finding that when combined with our work suggests that dihydroceramide accumulation in the ER is the proximal cause of HLD-18.

Although ER expansion represents the most proximal effect of loss of *ifc/DEGS1* function and dihydroceramide accumulation on cortex glia, other organelles and their functions are likely also disrupted, minimally as a consequence of ER disruption. For example, the apparent reduction of the lysosome marker, LAMP, in cortex glia of *ifc* mutant larvae correlates with increased RFP levels and the presence of bright RFP puncta or aggregates in this cell type, suggesting impaired lysosome function contributes to increased RFP perdurance and aggregation. Defects in the activity of multiple organelles may collaborate to elicit the full phenotype manifested by loss of *ifc/DEGS1* function, resulting in glial dysfunction and subsequent neuronal cell death.

Increased dihydroceramide levels may contribute to a broader spectrum of neurodegenerative diseases than simply HLD-18. Recent work reveals that gain of function mutations in *SPTLC1* and *SPTLC2*, which encode components of the SPT complex that catalyzes the initial, rate limiting step of *de novo* ceramide and sphingolipid biosynthesis,^55^ cause juvenile ALS via increased sphingolipid biosynthesis.^18–22^ Of all sphingolipids, the relative levels of dihydroceramide were increased the most in patient plasma samples, suggesting that DEGS1 activity becomes limiting in the presence of enhanced SPT activity and that dihydroceramide accumulation contributes to juvenile ALS. Any mutations that increase metabolite flux through the ceramide pathway upstream of DEGS1 may then increase dihydroceramide levels, drive ER expansion and cell swelling, and lead to neurodegeneration, with disease severity predicated on the extent of excessive dihydroceramide accumulation. Model systems, like flies, can harness the power of genetic modifier screens to identify genes and pathways (potential therapeutic targets), that can be tweaked to ameliorate the effect of elevated dihydroceramide levels on neurodegeneration.

Our work appears to pinpoint glia as the cell type impacted by loss of *ifc/DEGS1* function, specifically glia that exhibit great demand for membrane biogenesis like cortex glia in the fly and oligodendrocytes or Schwann cells in mammals. In larvae, *ifc* is expressed at higher levels in glial cells than in neurons, and its genetic function is required at a greater level in glia than neurons to govern CNS development. Our unpublished work on other genes in the ceramide metabolic pathway reveals similar glial-centric expression patterns in the larval CNS to that observed for *ifc*, suggesting they too function primarily in glia rather than neurons at this stage. In support of this model, a study on CPES, which converts ceramide into CPE, the fly analog of sphingomyelin, revealed that CPES is required in cortex glia to promote their morphology and homeostasis and to protect flies from photosensitive epilepsy.^59^ Similarly, ORMDL, a dedicated negative regulator of the SPT complex, is required in oligodendrocytes to maintain proper myelination in mice.^60^ Given that many glial cell types are enriched in sphingolipids and exhibit a great demand for new membrane biogenesis during phases of rapid neurogenesis and axonogenesis, we believe that glial cells, such as cortex glia, oligodendrocytes, and Schwann cells, rather than neurons manifest a greater need for *ifc/DEGS1* function and ceramide synthesis during developmental stages marked by significant nervous system growth.

Will this glial-centric model for *ifc/DEGS1* function, and more generally ceramide synthesis, hold true in the adult when neurogenesis is largely complete and the demand for new membrane synthesis in glia dissipates? Recent work in the adult fly eye suggests it may not. Wang et al.^61^ argued that the GlcT enzyme, which converts ceramide to glucosylceramide, is expressed at much higher levels in neurons than glia, and that glucosylceramide is then transported from neurons to glial cells for its degradation, suggesting cell-type specific compartmentalization of sphingolipid synthesis in neurons and its degradation in glia in the adult. In the future, it will be exciting to uncover whether genes of the sphingolipid metabolic pathway alter their cell-type specific requirements as a function of developmental stage.

We note that cortex glia are the major phagocytic cell of the CNS and phagocytose neurons targeted for apoptosis as part of the normal developmental process.^23–26^ Thus, while we favor the model that *ifc* triggers neuronal cell death due to glial dysfunction, it is also possible that increased detection of dying neurons arises due at least in part to a decreased ability of cortex glia to clear dying neurons from the CNS. At present, the large number of neurons that undergo developmentally programmed cell death combined with the significant disruption to brain and ventral nerve cord morphology caused by loss of *ifc* function render this question difficult to address. Additional evidence does, however, support the idea that loss of *ifc* function drives excess neuronal cell death: Clonal analysis in the fly eye reveals that loss of *ifc* drives photoreceptor neuron degeneration^17^, indicating that loss of *ifc* function drives neuronal cell death; cortex-glia specific depletion of *CPES*, which acts downstream of *ifc*, disrupts neuronal function and induces photosensitive epilepsy in flies^59^, indicating that genes in the ceramide pathway can act non-autonomously in glia to regulate neuronal function; recent genetic studies reveal that other glial cells can compensate for impaired cortex glial cell function by phagocytosing dying neurons^62^, and we observe that the cell membranes of subperineurial glia enwrap dying neurons in *ifc* mutant larvae (Fig. S14), consistent with similar compensation occurring in this background, and in humans, loss of function mutations in *DEGS1* cause neurodegeneration.^7–9^ Clearly, future work is required to address this question for *ifc/DEGS1* and perhaps other members of the ceramide biogenesis pathway.

Altered glial function may not only derive from dihydroceramide buildup in the ER, but also from altered cell membrane structure due to the replacement of ceramide and its derivates, such as GlcCer and CPE, with the cognate forms of dihydroceramide. Relative to dihydroceramide species, the 4-5 *trans* carbon-carbon double bond in the sphingoid backbone of ceramide-containing sphingolipids enables them to form more stable hydrogen bonds with water molecules and facilitates their ability to associate with lipids of different saturation levels.^5^ A high dihydroceramide to ceramide ratio has been shown to form rigid gel-like domains within model membranes and to destabilize biological membranes by promoting their permeabilization.^63,64^ As even minor alterations to membrane properties can disrupt glial morphology,^65^ such alterations in membrane rigidity and stability may underlie the failure of cortex glia to enwrap adjacent neurons. The observed increase in saturation of PE, PC, and PS in the CNS of *ifc* mutant larvae may reflect a compensatory response employed by cells to stabilize cell membranes and guard cell integrity when challenged with elevated levels of dihydroceramide.

The expansion of ER membranes coupled with loss of lipid droplets in *ifc* mutant larvae suggests that the apparent demand for increased membrane phospholipid synthesis may drive lipid droplet depletion, as lipid droplet catabolism can release free fatty acids to serve as substrates for lipid synthesis. At some point, the depletion of lipid droplets, and perhaps free fatty acids as well, would be expected to exhaust the ability of cortex glia to produce additional membrane phospholipids required for fully enwrapping neuronal cell bodies. Under wild-type conditions, many lipid droplets are present in cortex glia during the rapid phase of neurogenesis that occurs in larvae. During this phase, lipid droplets likely support the ability of cortex glia to generate large quantities of membrane lipids to drive membrane growth needed to ensheath newly born neurons. Supporting this idea, lipid droplets disappear in the adult *Drosophila* CNS when neurogenesis is complete and cortex glia remodeling stops.^30^ We speculate that lipid droplet loss in *ifc* mutant larvae contributes to the inability of cortex glia to enwrap neuronal cell bodies. Prior work on lipid droplets in flies has focused on stress-induced lipid droplets generated in glia and their protective or deleterious roles in the nervous system.^66–68^ Work in mice and humans has found that more lipid droplets are often associated with the pathogenesis of neurodegenerative diseases,^69–71^ but our work correlates lipid droplet loss with CNS defects. In the future, it will be important to determine how lipid droplets impact nervous system development and disease.

## Acknowledgements

We thank the Iowa Developmental Studies Hybridoma Bank for antibodies, and the Bloomington Stock Center, Vienna Drosophila Research Center, FlyORF, and the National Institutes of Genetics stock center in Japan for countless fly lines. We thank Drs. Christian Klambt, Chih-Chiang Chan, Dion Dickman for reagents. We thank Dr. Tristan Qingyuan Li for comments on the manuscript. We thank the Genome Technology Access Center at Washington University for next-generation sequencing and analysis of RNA-seq samples. We thank Dr. Sanja Sviben, Gregory Strout, and John Wulf II for assistance in TEM studies conducted at the Washington University Center for Cellular Imaging, which is supported in part by Washington University School of Medicine, The Children’s Discovery Institute of Washington University and St. Louis Children’s Hospital (CDI-CORE-2015-505 and CDI-CORE-2019-813), the Foundation for Barnes-Jewish Hospital (3770) and the Washington University Diabetes Research Center (NIH P30 DK020579).

## Author Contributions

BAW, HL, Yi Zhu, and JBS completed the genetic screen that identified the *ifc* alleles and completed the initial genetic characterization of these alleles. Yuqing Zhu, JD, and JBS completed all other experiments except for generating the lipidomics data via mass spectrometry, which were completed by KC and GJP. Yuqing Zhu and JBS wrote the manuscript and compiled figures with help from JD with figures. All authors read, commented on, and edited the manuscript.

## Methods

### Key resources table

#### Fly stocks used

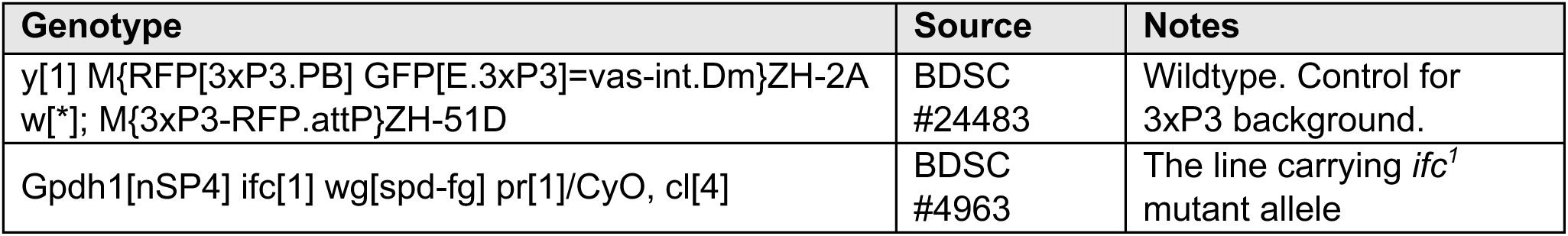
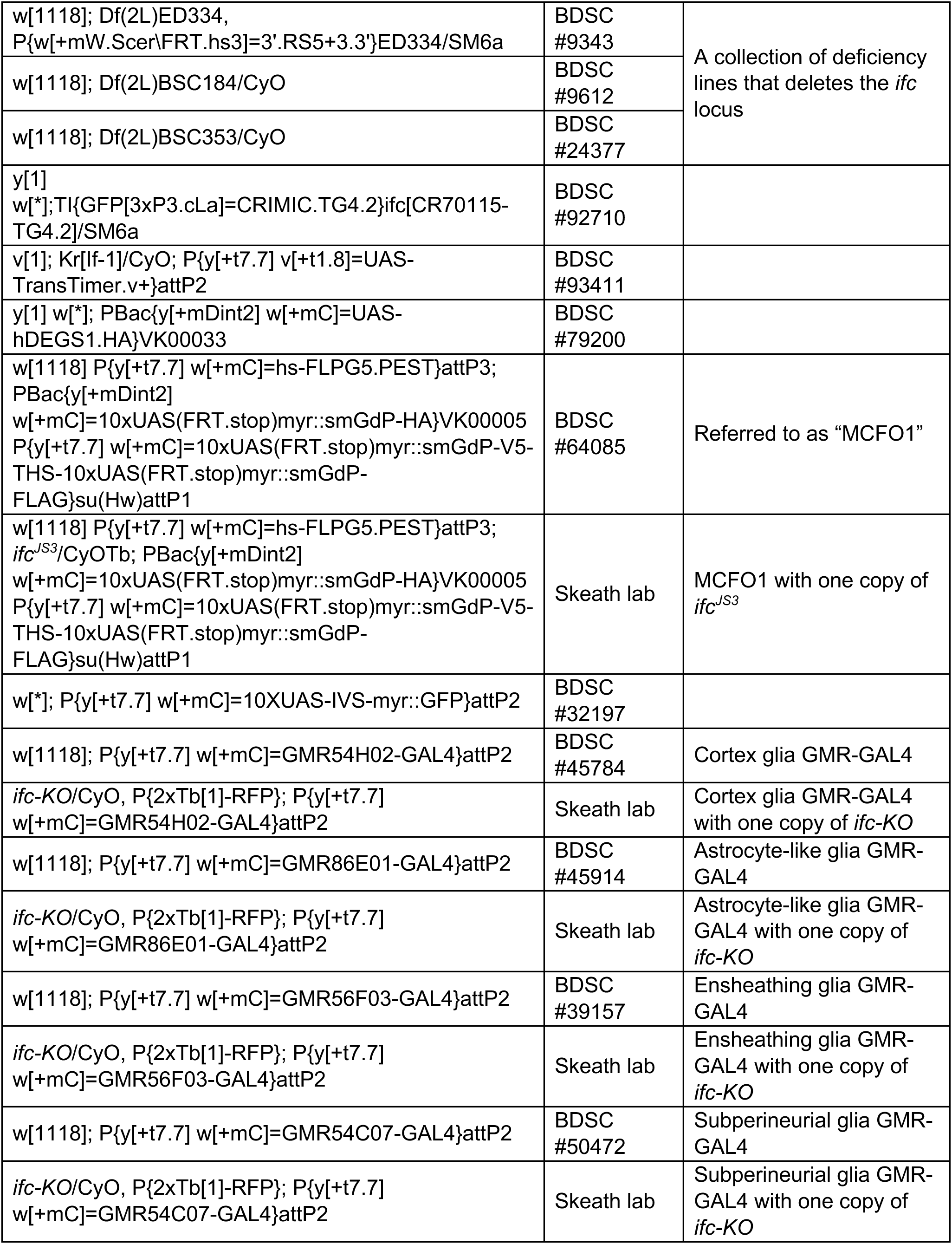
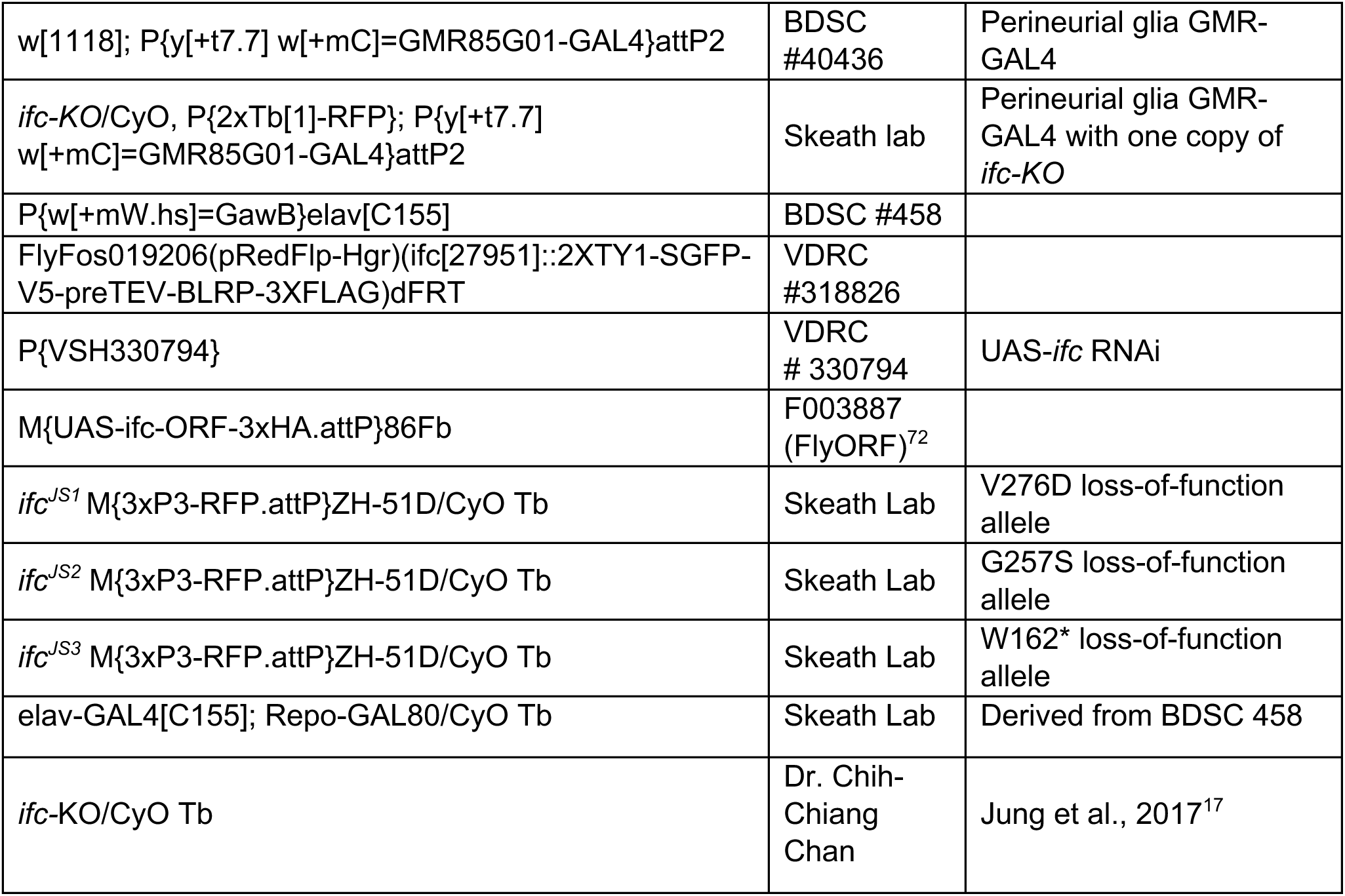

#### Antibodies and lipid marker used

| **Antibody/marker** | **Dilution** | **Source** | **RRID** |
| --- | --- | --- | --- |
| Rabbit anti-FABP | 1:500 | Skeath lab |  |
| Rabbit anti-EBONY | 1:500 | Gift from Dr. Haluk Lacin | AB_2314354 |
| Mouse anti-REPO | 1:100 | DSHB | AB_528448 |
| Rat anti-ELAV | 1:100 | DSHB | AB_528218 |
| Mouse anti-CXN99A | 1:20 | DSHB | AB_2722011 |
| Mouse anti-GOLGIN84 | 1:20 | DSHB | AB_2722113 |
| Goat anti-GOLGIN245 | 1:500 | DSHB | AB_2618260 |
| Rabbit anti-ESYT | 1:300 | Gift from Dr. Dion Dickman |  |
| GFP Antibody Dylight™ 488 Conjugated Pre-Adsorbed, Goat Polyclonal | 1:500 | Rockland (600-141-215) | AB_1961516 |
| Anti-LAMP1 antibody - Lysosome Marker | 1:500 | Abcam (ab 30687) | AB_775973 |
| Cleaved Caspase-3 (Asp175) Antibody | 1:400 | Cell Signaling Technology® (#9661) | AB_2341188 |
| Anti-HA antibody produced in rabbit | 1:500 | Millipore Sigma (H6908) | AB_260070 |
| ANTI-FLAG® antibody, Rat monoclonal | 1:500 | Millipore Sigma (SAB4200071) | AB_10603396 |
| Chicken V5 Tag Polyclonal Antibody | 1:500 | Bethyl laboratories (A190-118A) | AB_66741 |
| BODIPY™ 493/503 (4,4-Difluoro-1,3,5,7,8-Pentamethyl-4-Bora-3a,4a-Diaza-s-Indacene) | 1:200 | Invitrogen™ (D3922) |  |
| Goat anti-Chicken IgY (H+L) Secondary Antibody, Alexa Fluor™ 488 | 1:1000 | Invitrogen™ (A-11039) | AB_2534096 |
| Donkey anti-Rat IgG (H+L) Highly Cross-Adsorbed Secondary Antibody, Alexa Fluor™ Plus 555 | 1:1000 | Invitrogen™ (A48270) | AB_2896336 |
| Cy™5 AffiniPure Donkey Anti-Rat IgG (H+L) | 1:1000 | Jackson ImmunoResearch (712-175-153) | AB_2340672 |
| Cy™5 AffiniPure Donkey Anti-Goat IgG (H+L) | 1:1000 | Jackson ImmunoResearch (705-175-147) | AB_2340415 |
| Cy™5 AffiniPure Donkey Anti-Mouse IgG (H+L) | 1:1000 | Jackson ImmunoResearch (715-175-151) | AB_2340820 |
| Cy™5 AffiniPure Donkey Anti-Rabbit IgG (H+L) | 1:1000 | Jackson ImmunoResearch (711-175-152) | AB_2340607 |

#### Deposited data

|  |  |  |
| --- | --- | --- |
| Raw and processed RNA sequencing data | GEO: GSE263308 | https://www.ncbi.nlm.nih.gov/geo/query/acc.cgi?acc=GSE263308 |
| Untargeted lipidomics | PR001967: ST003162 | https://dx.doi.org/10.21228/M8G72D |

#### Software used

| **Software** | **Source** | **Website** |
| --- | --- | --- |
| ImageJ2 2.3.0/1.53q | NIH; Schindelin et al., 2012 | https://imagej.net/ij/ |
| ZEN Microscopy Software | Carl Zeiss AG; Jena, Germany | https://www.zeiss.com/microscopy/en/home.html |
| Photoshop 23.5.5 | Adobe; San Jose, California, USA | https://www.adobe.com/products/photoshop.html |
| GraphPad Prism 10.2.2 | GraphPad; Boston, MA, USA | https://www.graphpad.com/features |
| BioRender | BioRender; Toronto, CAN | https://www.biorender.com |
| Microsoft Office 365 16.83 (24031120) | Microsoft; Redmond, USA | https://www.microsoft.com/en-us/microsoft-365 |

## Method details

### Fly husbandry

Flies were raised on standard molasses-based food at 25°C. Unless otherwise noted, wild type is an otherwise wild-type stock harboring the *M{3xP3-RFP.attP}ZH-51D* insert in an isogenic second chromosome.

### Mutagenesis

A standard autosomal recessive forward genetic screen was carried out using 25-30 μm EMS to mutagenize a *M{3xP3-RFP.attP}ZH-51D* isogenic second chromosome. Homozygous mutant third instar larvae were visually screened under a standard fluorescent microscope for defects in CNS morphology.A detailed description of the screen, all identified genes, and associated whole genome sequences are described elsewhere36.

### Creation of recombinant lines and identification of second site mutations in *ifc-KO* **chromosome**

Standard genetic methods were used to generate fly strains that contained specific combinations of GAL4 and UAS-linked transgenes in the *ifc^js3^*or *ifc-KO* background. During this process, we uncovered that the *ifc-KO* deletion could be unlinked from at least one second chromosomal mutation that caused early larval lethality, resulting in homozygous *ifc-KO* flies that survived to late L3 to early pupa. A subsequent EMS-based F2 lethal non-complementation screen using the *M{3xP3-RFP.attP}ZH-51D* isogenic second chromosome as target chromosome and screening against the *ifc-KO* chromosome identified multiple mutations in four complementation groups that led to an early larval lethal phenotype. Whole genome sequencing identified these genes as *Med15*, *lwr*, *Nle*, and *Sf3b1*; all four genes reside within ∼90 Kb of each other in chromosomal bands 21D1-21E2 near the telomere in chromosome 2L, identifying the site of the associated lethal mutations and suggesting the actual lesion may be a small deletion that removes these genes. The following alleles of these genes are available at Bloomington Stock Center: *Med15^js1^*(Q175*), *Med15^js2^* (Q398*), *Med15^js3^* (C655Y), *lwr^js1^* (I4N), *lwr^js2^*(E12K), *Nle^js1^*(W146R), *Nle^js2^* (L125P), *Nle^js3^* (Q242*), *Sf3b1^js1^* (R1160*), *Sf3b1^js^*^2^ (Q1264*), *Sf3b1^js3^* (570 bp deletion at the following coordinates chr2L571720-572290; this deletion removes amino acids 1031 through 1222 and introduces a frameshift into the reading frame).

### Gene rescue and in vivo RNAi phenocopy assays

To restrict UAS-linked transgene expression specifically to glia, we used the *repoGAL4* driver line. To restrict UAS-linked transgene expression specifically to neurons, we paired the *elavGAL4* driver lines, which activates transgene expression strongly in all neurons and moderately in glia, with *repoGAL80*, which blocks GAL4-dependent activation in glia. We used *P{VSH330794}* (VDRC 330794) for the RNAi experiments and *UAS-ifc*^72^ and *UAS-DEGS1* (BDSC 79200) for the gene rescue assays. All gene rescue experiments were performed in the *ifc^js3^*/*ifc-KO* background with the UAS-transgene placed into the *ifc^js3^* background and the GAL4 drivers into the *ifc-KO* background.

The GAL4-UAS method was also used to assess the phenotype of each glial subtype. In combination with the *UAS-Myr-GFP* transgene, which labels cell membranes, we used the following GAL4 lines to trace the morphology of each glial subtype in wild type and *ifc^-/-^* larvae: *GMR85G01-GAL4* (perineurial glia; BDSC 40436), *GMR54C07-GAL4* (subperineurial glia; BDSC 50472), *GMR54H02-GAL4* (cortex glia; BDSC 45784), *GMR56F03-GAL4* (ensheathing glia; BDSC 77469 and 39157), and *GMR86E01-GAL4* (astrocyte-like glia; BDSC45914). The *UAS-Myr-GFP* transgene was placed into the *ifc^js3^*background; each glial GAL4 line was placed into the *ifc-KO* background.

The GAL4-UAS method was used to assess the ability of *UAS-ifc* and *UAS-DEGS1* transgenes to rescue the lethality of otherwise *ifc* mutant larvae. *ifc-KO/CyO Tb; repo-GAL4/TM6B Tb* males were crossed to each of the following lines: *ifc^js3^/CyO Tb; UAS-ifc*; *ifc^js3^/CyO Tb; UAS-DEGS1*; *ifc^js3^/CyO Tb.* All adult flies were sorted into Curly and non-Curly and then counted; all non-Curly flies lacked the *TM6B Tb* balancer indicating they carried *repo-GAL4* and thus were likely rescued to viability by glial expression of *ifc* or *DEGS1*. Standard Mendelian ratios were then used to predict the expected number of *ifc* mutant flies if all survived to adulthood. The total number of observed adult *ifc* mutant flies, identified by lack of Curly wings, was then divided by this number to obtain the percentage of *ifc* mutant flies that survived to adulthood. 2452 total flies were assayed for the *UAS-ifc* cross, 1303 for the *UAS-DEGS1* cross, and 1030 for the control cross.

### DNA sequencing

Genomic DNA was obtained from wild type larvae or larvae homozygous for each relevant mutant line and provided to GTAC (Washington University) or GENEWIZ for next-generation or Sanger sequencing.

### RNA in-situ hybridization

*ifc* RNA probes for in-situ hybridization chain reaction (HCR) were designed and made by Molecular Instruments (HCR™ RNA-FISH v3.0).^73^ Wild type CNS was harvested and fixed in 2% paraformaldehyde at late L3. The fixed CNS underwent gradual dehydration and rehydration, followed by standard hybridization and amplification steps of the HCR protocol.^73^ For double labeling with antibody, the post-HCR labeled CNS was briefly fixed for 30 minutes prior to standard antibody labeling protocol to identify specific CNS cell type(s) with highly localized *ifc* _RNAs.74,75_

### MultiColor FlpOut labeling of glial subtypes

For glial labeling in the control background, the *MCFO1* line was crossed to *GMR-GAL4* driver lines for each glial subtype (Supplemental Table 1).^33^ For glial labeling in *ifc^-/-^* background, the *MCFO1* line was placed in the *ifc^JS3^* background, each of the five glial-specific GMR-GAL4 lines were placed individually into the *ifc-KO* background, and then the MCFO, *ifc^JS3^* line was crossed to each glial specific GAL4, *ifc-KO* line. Flies were allowed to lay eggs for 24 hr at 25°C, and progeny were raised at 25°C for 4 days prior to heat-activated labeling. On day 4 after egg-laying, F1 larvae were incubated in a 37°C water bath for 5-minutes. When wild-type or *ifc^-/-^* mutant larvae reached late L3, which was day 5-6 for control and day 9-10 for *ifc^-/-^* mutants, the CNS was dissected, fixed, stained, and then analyzed under a Zeiss LSM 700 Confocal Microscope for the presence of clones, using Zen software.

### Antibody generation

YenZym (CA, USA) was used as a commercial source to generate affinity purified antibodies against a synthetic peptide that corresponded to amino acids 85-100 (TLDGNKLTQEQKGDKP) of FABP isoform B. Briefly, the peptide was conjugated to KLH, used as an immunogen in rabbits to generate a peptide-specific antibody response, and antibodies specific to the peptide were affinity purified. The affinity-purified antibodies were confirmed to label cortex glia specifically based on comparison of antibody staining relative to Myr-GFP when a UAS-Myr-GFP was driven under control of the cortex specific glial GAL4 driver GMR-54H02 (Figure S3). The FABP antibodies are used at 1:500-1:1000.

### Immunofluorescence and lipid droplet staining

Gene expression analysis was performed essentially as described in Patel.^76^ Briefly, the larval CNS was dissected in PBS, fixed in 2.5% paraformaldehyde for 55 minutes, and washed in PTx (1xPBS, 0.1% TritonX-100. The fixed CNS was incubated in primary antibody with gentle rocking overnight at 4°C. Secondary antibody staining was conducted for at least two hours to overnight at room temperature. All samples were washed in PTx at least five times and rocked for an hour before and after secondary antibody staining. A detailed list of the primary and secondary antibodies is available in Supplemental Table 2. Dissected CNS were mounted either in PTx or dehydrated through an ethanol series and cleared in xylenes prior to mounting in DPX mountant.^77^ All imaging was performed on a Zeiss LSM-700 Confocal Microscope, using Zen software.

For lipid droplet staining, the fixed CNS was incubated for 30 minutes at room temperature at 1:200 dilution of 1mg/mL BODIPY™ 493/503 (Invitrogen: D3922). It was then rinsed thoroughly in PBS and immediately mounted for imaging on a Zeiss LSM-700 Confocal Microscope, using Zen software.

### Transmission electron microscopy (TEM) imaging

For transmission electron microscopy (TEM), samples were immersion fixed overnight at 4°C in a solution containing 2% paraformaldehyde and 2.5% glutaraldehyde in 0.15 M cacodylate buffer with 2 mM CaCl2, pH 7.4. Samples were then rinsed in cacodylate buffer 3 times for 10 minutes each and subjected to a secondary fixation for one hour in 2% osmium tetroxide/1.5% potassium ferrocyanide in cacodylate buffer. Following this, samples were rinsed in ultrapure water 3 times for 10 minutes each and stained overnight in an aqueous solution of 1% uranyl acetate at 4°C. After staining was complete, samples were washed in ultrapure water 3 times for 10 minutes each, dehydrated in a graded acetone series (50%, 70%, 90%, 100% x4) for 15 minutes in each step, and infiltrated with microwave assistance (Pelco BioWave Pro, Redding, CA) into Spurr’s resin. Samples were then cured in an oven at 60°C for 72 hours and post-curing, 70 nm thin sections were cut from the resin block, post-stained with uranyl acetate and Sato’s lead and imaged on a Transmission Electron Microscope (JEOL JEM-1400 Plus, Tokyo, Japan) operating at 120 KeV.

### Lipidomics

Untargeted lipidomics analysis was conducted on whole larva and dissected CNS of wild type and *ifc^-/-^* mutants at the late-third instar stage. Five replicates were prepared for each set of experiments. For whole larvae, at least 15 larvae of each genotype were used for each replicate. For the dissected CNS, at least 50 wild type and 60 *ifc*^-/-^ CNS were used per replicate. Immediately following collection or dissection, larvae and the dissected CNS were flash frozen in liquid nitrogen and placed at −80°C.

Lipids were extracted from frozen whole larvae and dissected larval CNS samples by using an Omni Bead Ruptor Elite Homogenizer using acetonitrile:methanol:water (2:2:1; 40 μL/mg tissue). Two Ultrahigh performance LC-(UHPLC)/MS systems were used in this work: a Thermo Vanquish Flex UHPLC system with a Thermo Scientific Orbitrap ID-X and an Agilent 1290 Infinity II UPLC system with an Agilent 6545 QTOF as described previously.^78^ Lipids were separated on a Waters Acquity HSS T3 column (150 x 2.1 mm, 1.8 mm). The mobile-phase solvents were composed of A: 0.1% formic acid, 10 mM ammonium formate, 2.5 μM medronic acid in 60:40 acetonitrile:water; and B = 0.1% formic acid, 10 mM ammonium formate in 90:10 2-propanol:acetonitrile. The column compartment was maintained at 60 °C. The following linear gradient was applied at a flow rate of 0.25 mL min-1: 0 – 2 min, 30% B; 17 min, 75% B; 20 min, 85% B; 23 – 26 min, 100% B. The injection volume 4 μL for all lipids analysis. Data was acquired in positive ion.

LC/MS data were processed and analyzed with the open-source Skyline software.^79^ Lipid MS/MS data were annotated with Agilent Lipid Annotator software.

### RNA sequencing and analysis

To determine the CNS-specific transcriptional changes upon loss of *ifc*, RNA-seq was conducted on five replicates of dissected CNS tissue derived from wild type and *ifc^-/-^* mutant late-third instar larvae. For each replicate, roughly 30-35 dissected CNS of wild type or *ifc-/-* larvae were used. Invitrogen™ RNAqueous™-Micro Total RNA Isolation Kit (AM1931) was used to extract RNA. Agilent 4200 TapeStation system was used for RNA quality control.

Samples were prepared according to library kit manufacturer’s protocol, indexed, pooled, and sequenced on an Illumina NovoSeq. Basecalls and demultiplexing were performed with Illumina’s bcl2fastq software and a custom python demultiplexing program with a maximum of one mismatch in the indexing read. RNA-seq reads were then aligned to the Ensembl release 76 primary assembly with STAR version 2.5.1a.^80^ Gene counts were derived from the number of uniquely aligned unambiguous reads by Subread:featureCount version 1.4.6-p5.^81^ Isoform expression of known Ensembl transcripts were estimated with Salmon version 0.8.2.^82^ To find the most critical genes, the raw counts were variance stabilized with the R/Bioconductor package DESeq2^83^ and was then analyzed via weighted gene correlation network analysis with the R/Bioconductor package WGCNA.^84^

### Statistics

All data are presented as mean ± SEM. Statistical significance between groups was determined using Student’s *t*-test or one-way ANOVA with multiple comparisons, and with varying levels of significance assessed as * *P* < 0.05, ** *P* < 0.01, *** *P* < 0.001, **** *P* < 0.0001, and ns, not significant.

## Supplemental Figures and information

**Figure S1.**
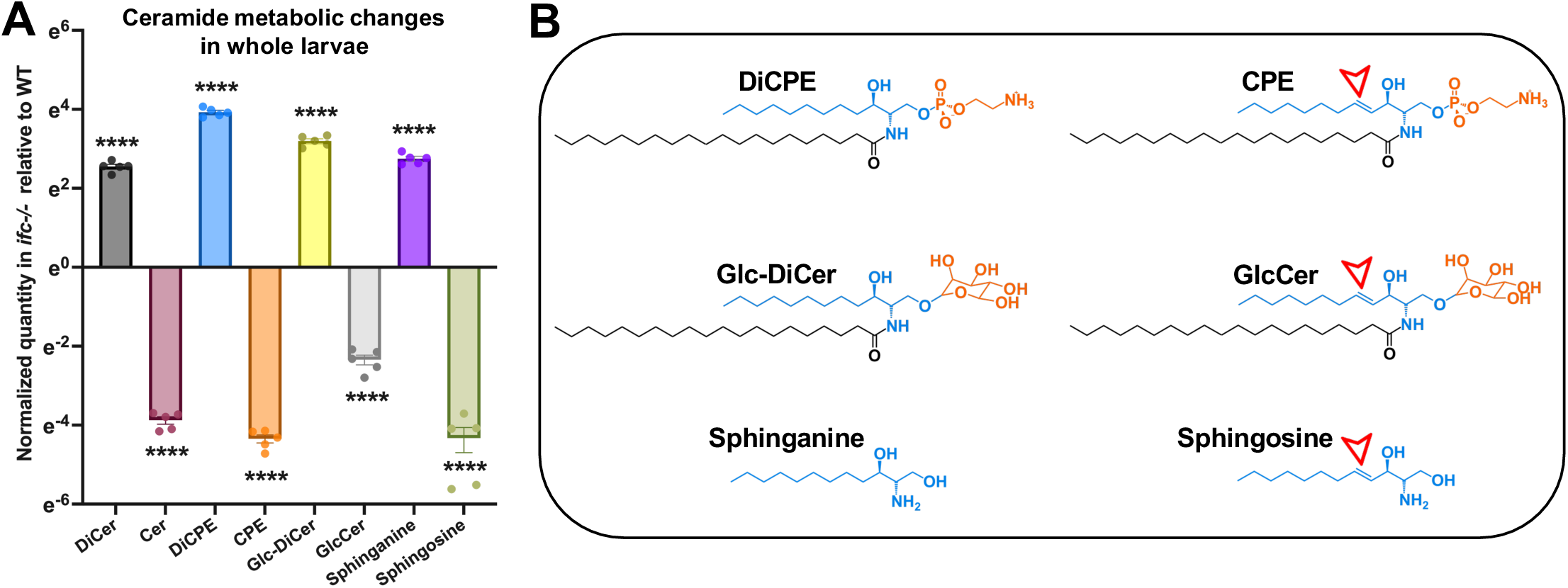
Ceramide metabolic changes in whole larvae. A) Normalized quantification of the relative levels of dihydroceramide, ceramide, and six related sphingolipid species in the whole larvae of wild-type and *ifc^-/-^* late-third instar larvae. B) Chemical structure of dihydroceramide phosphoethanolamine (DiCPE), ceramide phosphoethanolamine (CPE), glucosyl-dihydroceramide (Glc-DiCer), glucosyl-ceramide (GlcCer), sphinganine, and sphingosine; arrow indicates the cis carbon-carbon double bond between C4 and C5 in the sphingoid backbone created by the enzymatic action of Ifc/DEGS1.

**Figure S2).**
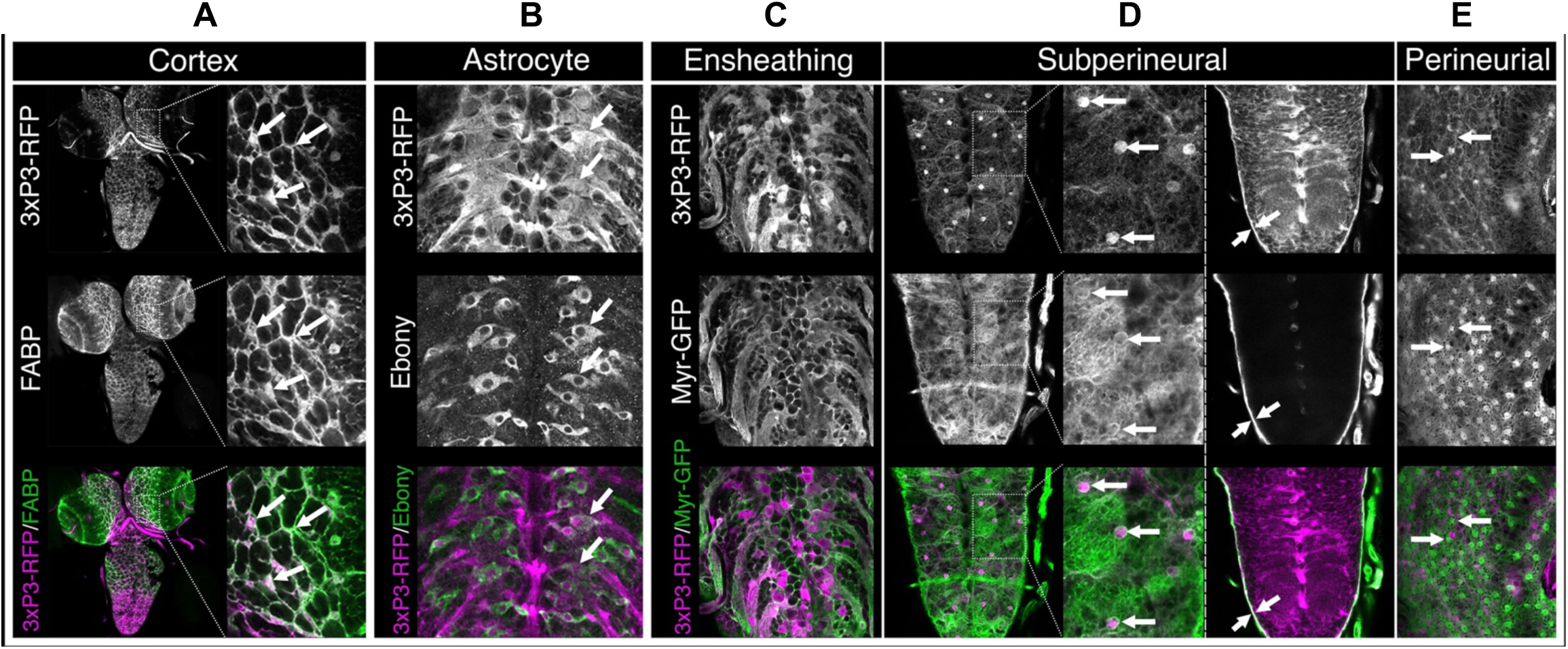
The *M{3xP3-RFP.attP}ZH-51D* transgene insert labels most glia. Low and high magnification images of late third instar larvae that harbor the *M{3xP3-RFP.attP}ZH-51D* transgene showing RFP expression and markers for different glial subtypes. **A) Cortex glia**: Ventral view of full CNS labeled for FABP to label cortex glia and RFP. Inset shows high magnification view of the brain with clear overlap of FABP and RFP expression (arrows). B) **Astrocyte-like glia**: Dorsal view of thoracic and abdominal region of CNS labeled for Ebony to label astrocyte glia and RFP; arrows highlight overlap of RFP and Ebony expression in some astrocyte-like glia. **C) Ensheathing glia**: Dorsal view of thoracic and abdominal region of CNS labeled for RFP and Myr-GFP. The *UAS-Myr-GFP* transgene is driven by the ensheathing glia-specific GAL4 driver line *GMR56F03*. Note the colocalization of RFP and GFP in the nerves exiting the CNS and the circular structures of ensheathing glia near the dorsal midline. **D) Subperineurial glia**: The two left-most panels show ventral views of thoracic and abdominal region of CNS labeled for RFP and Myr-GFP driven by the *GMR54C07-GAL4* line, which is specific to subperineurial glia. Arrows identify nuclei of subperineurial glia, which contain RFP and are encircled by Myr-GFP. Right panel shows cross section of CNS that highlights RFP and Myr-GFP co-expression in subperineurial glia at lateral edge of CNS (arrows). **E) Perineurial glia**: Ventral view of brain labeled for RFP and Myr-GFP driven by the *GMR85G01-GAL4* line, which is thought to be specific for perineurial glia. Arrows point to the small GFP-expressing perineurial glia, most of which do not express detectable levels of RFP. Visual inspection and quantification of the relative expression of RFP to each glial subtype marker in at least five brain-nerve cord complexes revealed that essentially all cortex, astrocyte, ensheathing, and subperineurial glia expressed RFP, but that astrocyte-like glia expressed RFP at much lower levels than the other three glial subtypes, and that most perineurial glia did not express RFP.

**Figure S3).**
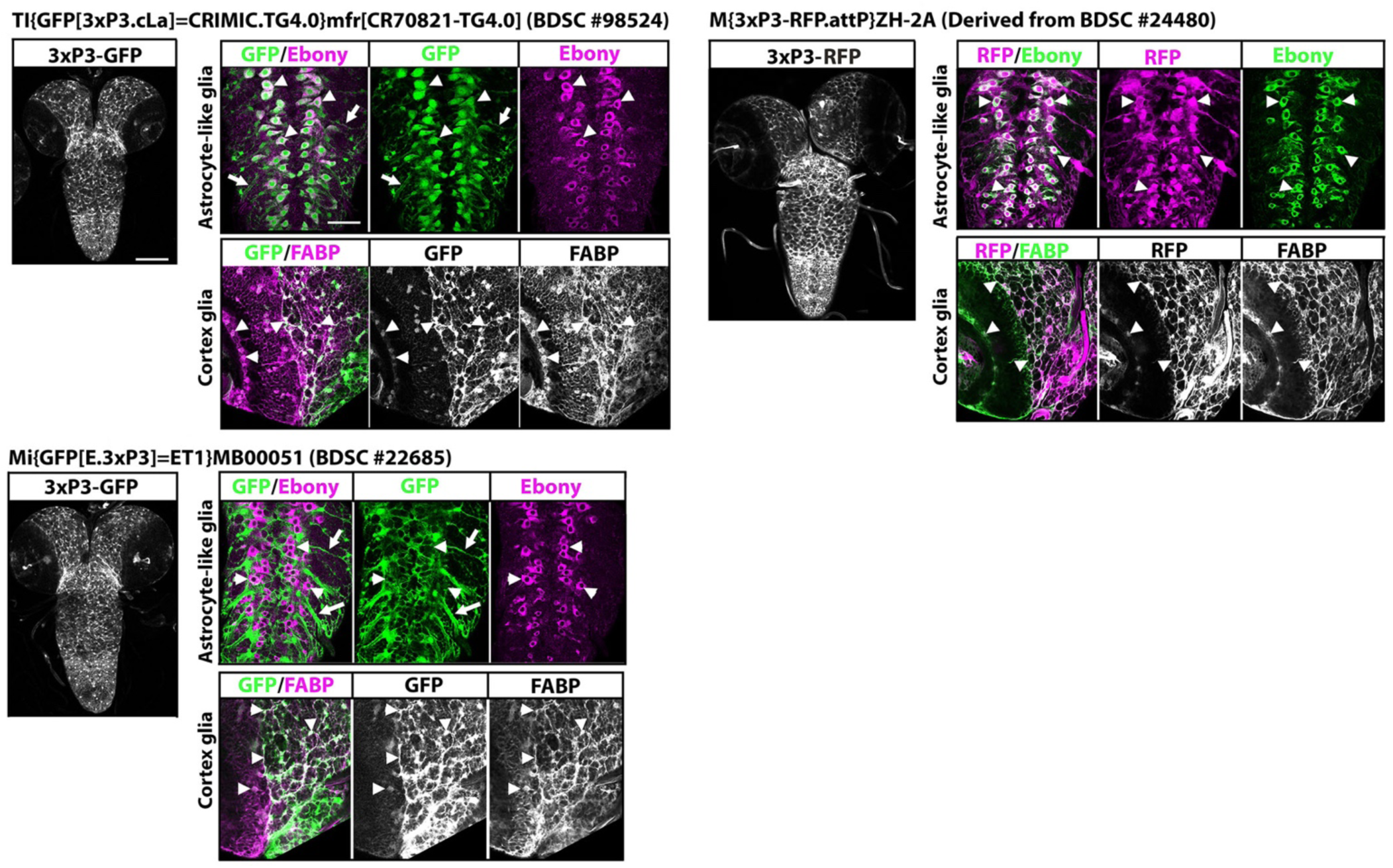
Most 3xP3-GFP or 3xP3-RFP transgenes label glia. Low magnification views of the indicated 3xP3 transgenes driving GFP (left) or RFP (right) in the CNS. Low magnification views illustrate that the three 3xP3 transgenes label or highlight the CNS. High magnification views illustrate that all three transgenes label cortex glia (arrowheads), which are marked by FABP. Two of the three transgenes strongly label astrocyte-like glia (arrowheads), which are marked by Ebony. Two of the three transgenes also clearly label ensheathing glia (arrows). Scale bar is 100 microns for low magnification images; 50 microns for high magnification images. We note that all M{3xP3-RFP.attp} transgenes (n=4), all Mi{GFP[E.3xP3]=ET1} transgenes (n=3), and all Tl{GFP[3xP3.cLa]=CRIMIC.TG4} transgenes (n=6) that we assayed labeled the CNS in a manner similar to that detailed here for the indicated lines. We note that we saw no GFP CNS expression in the TI{GFP[3xP3.cLa]=KozakGAL4}spag4[CR70688-KO-kG4]/SM6a line. All lines that we tested are listed below: **Mi{GFP[E.3xP3]=ET1} transgenes tested:** y[1] w[67c23]; Mi{GFP[E.3xP3]=ET1}MB00051: BDSC-22685 y[1] w[67c23]; Mi{GFP[E.3xP3]=ET1}MB00056: BDSC-22686 w[1118]; Mi{GFP[E.3xP3]=ET1}Ppn[MB07666]: BDSC 25277 **Tl{GFP[3xP3.cLa]=CRIMIC.TG4} transgenes tested:**y[1] w[*]; TI{GFP[3xP3.cLa]=CRIMIC.TG4.0}mfr[CR70821-TG4.0]: BDSC 98524: y[1] w[*]; TI{GFP[3xP3.cLa]=CRIMIC.TG4.0}lace[CR70094-TG4.0]/SM6: BDSC 92702 y[1] w[*] TI{GFP[3xP3.cLa]=CRIMIC.TG4.2}schlank[CR01622-TG4.2] Josd[CR01622-TG4.2-X]/FM7: BDSC 91266 y[1] w[*]; TI{GFP[3xP3.cLa]=CRIMIC.TG4.0}bwa[CR02807-TG4.0]/SM6a; BDSC 97620 y[1] w[*]; TI{GFP[3xP3.cLa]=CRIMIC.TG4.2}ifc[CR70115-TG4.2]/SM6a; BDSC 92710 y[1] w[*]; TI{GFP[3xP3.cLa]=CRIMIC.TG4.1}ghi[CR01805-TG4.1]/TM3; BDSC 92656 **M{3xP3-RFP.attp} transgenes tested** M{3xP3-RFP.attP}ZH-2A: Derived from BDSC 24480 M{3xP3-RFP.attP’}ZH-22A: Derived from BDSC 24481 M{3xP3-RFP.attP’}ZH-51C: Derived from BDSC 24482 {3xP3-RFP.attP}ZH-51D: Derived from BDSC 24483

**Figure S4).**
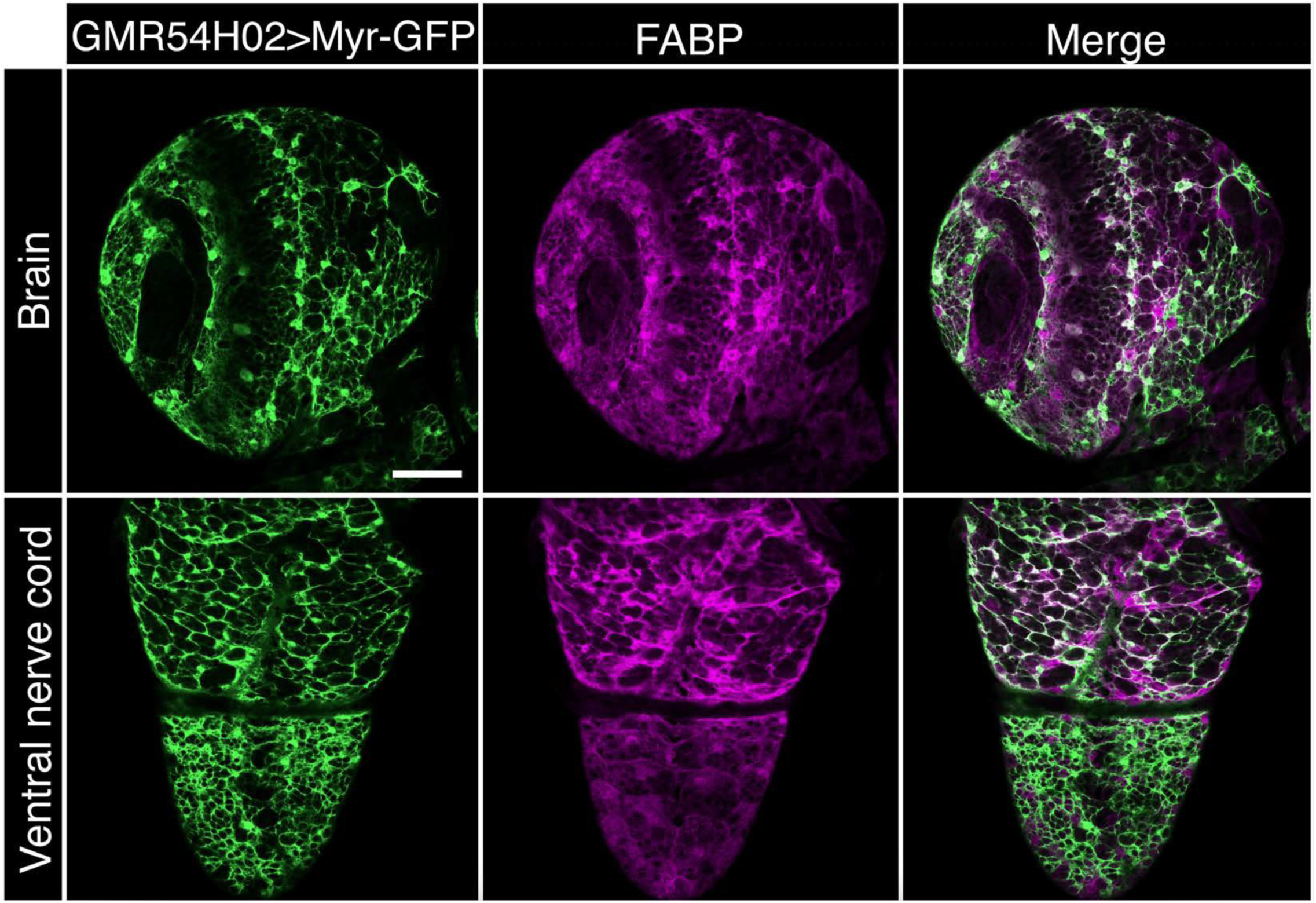
Newly generated FABP antibody labels cortex glia. Ventral views of the brain and ventral nerve cord of wild-type larvae that express Myr-GFP under the control of cortex-specific GAL4 driver line *GMR54H02*. Myr-GFP (green) highlights the cell membranes of cortex glia; FABP protein expression (red) exhibits significant overlap with Myr-GFP. Anterior is up; scale bar is 50 microns.

**Figure S5).**
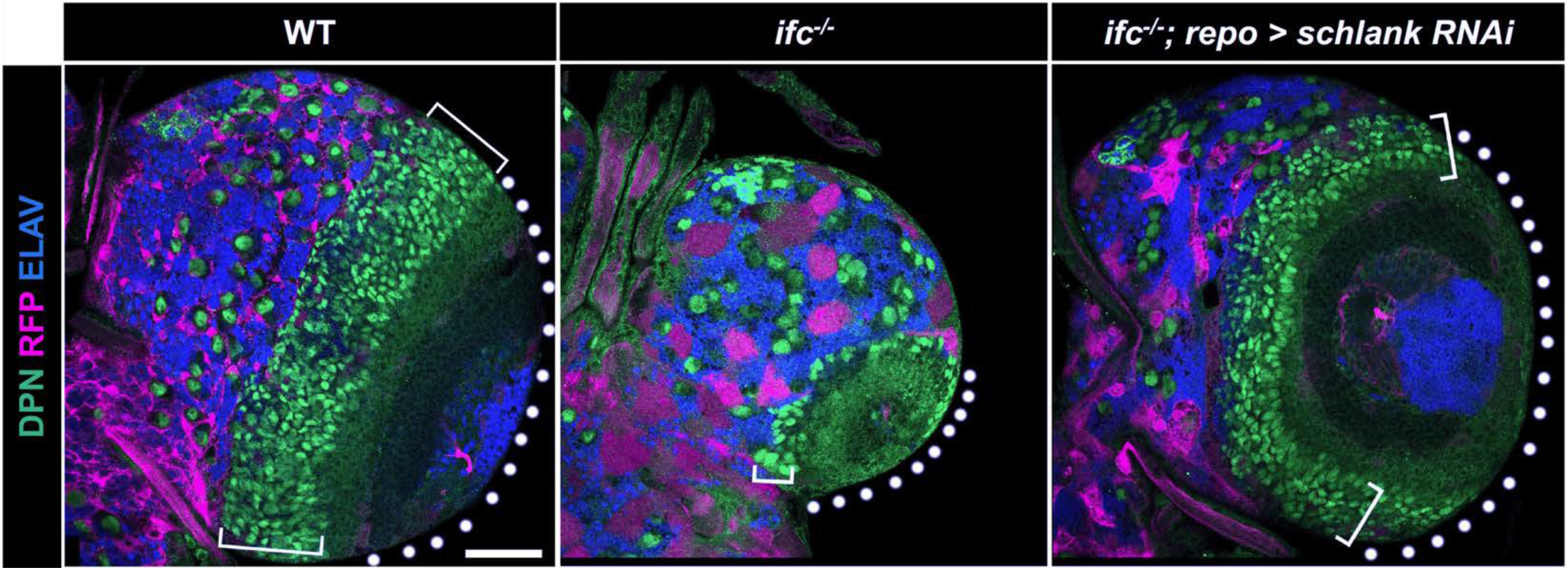
Loss of *ifc* function reduces optic lobe size and optic lobe neuroblast number. Ventral view of single brain hemisphere of late-third instar larvae of the indicated genotype labeled for neuroblasts (DPN; green), glia (RFP, red), and neurons (ELAV, blue). Loss of function in *ifc* results in reduction of the number of optic lobe neuroblasts (brackets) and the size of the optic lobe (dotted circles) relative to wild-type and *ifc* mutants in which *schlank* function is inhibited specifically in glia (*repo-GAL4>UAS-schlank^RNAi^).* Anterior is up; scale bar is 50 microns.

**Figure S6).**
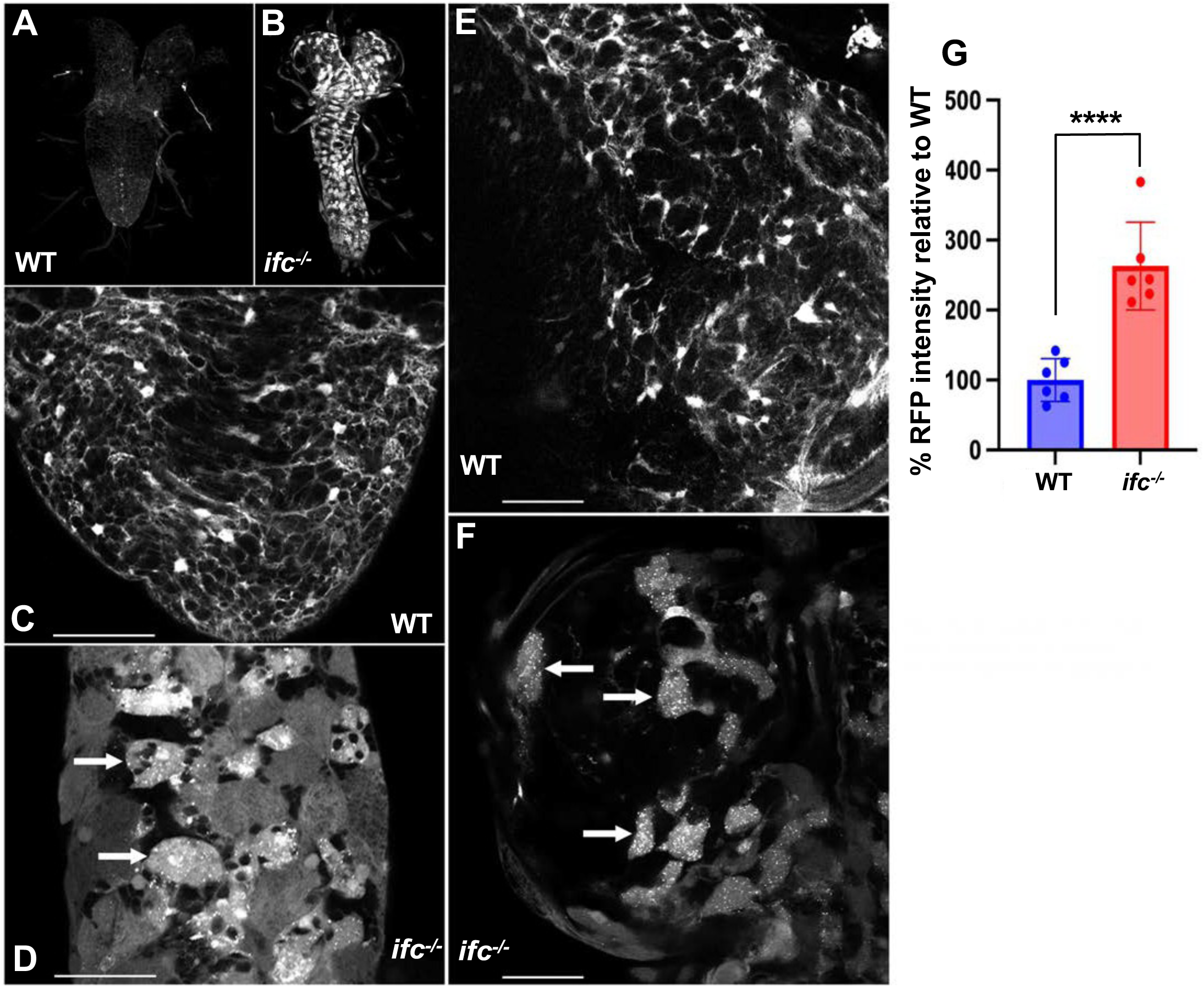
Loss of *ifc* increases RFP fluorescence and induces the formation of bright RFP-positive puncta or aggregates. A-B) Ventral views of the CNS of wildtype (A) and *ifc* mutant (B) late third instar larvae homozygous for the *M{3xP3-RFP.attP}* transgene, which drives RFP expression in most glia. RFP expression is shown in greyscale. Larvae were fixed at the same time in the same tube, mounted on the same slide, and imaged at the same time using identical parameters. *ifc* mutant larvae consistently exhibit higher levels of RFP expression. Anterior is up; scale bar is 100 microns. C-F) High magnification ventral views of the abdominal region of the ventral nerve cord (C and D) and brain (E and F) of wildtype (C and E) and *ifc* mutant larvae (D and F) showing RFP expression in greyscale. Note that the gain was increased in wildtype larvae (C and E) relative to *ifc* mutant larvae (D and F) to enable visualization of RFP signal in the wildtype CNS. Brightly fluorescent puncta or aggregates (arrows) appear in cortex glia of *ifc* mutant, but not wild-type larvae. Anterior is up. Scale bar is 50 microns for C-F. G) Quantification of relative RFP intensity between wild type and *ifc* mutant larvae – p<2.0×10^-4^.

**Figure S7).**
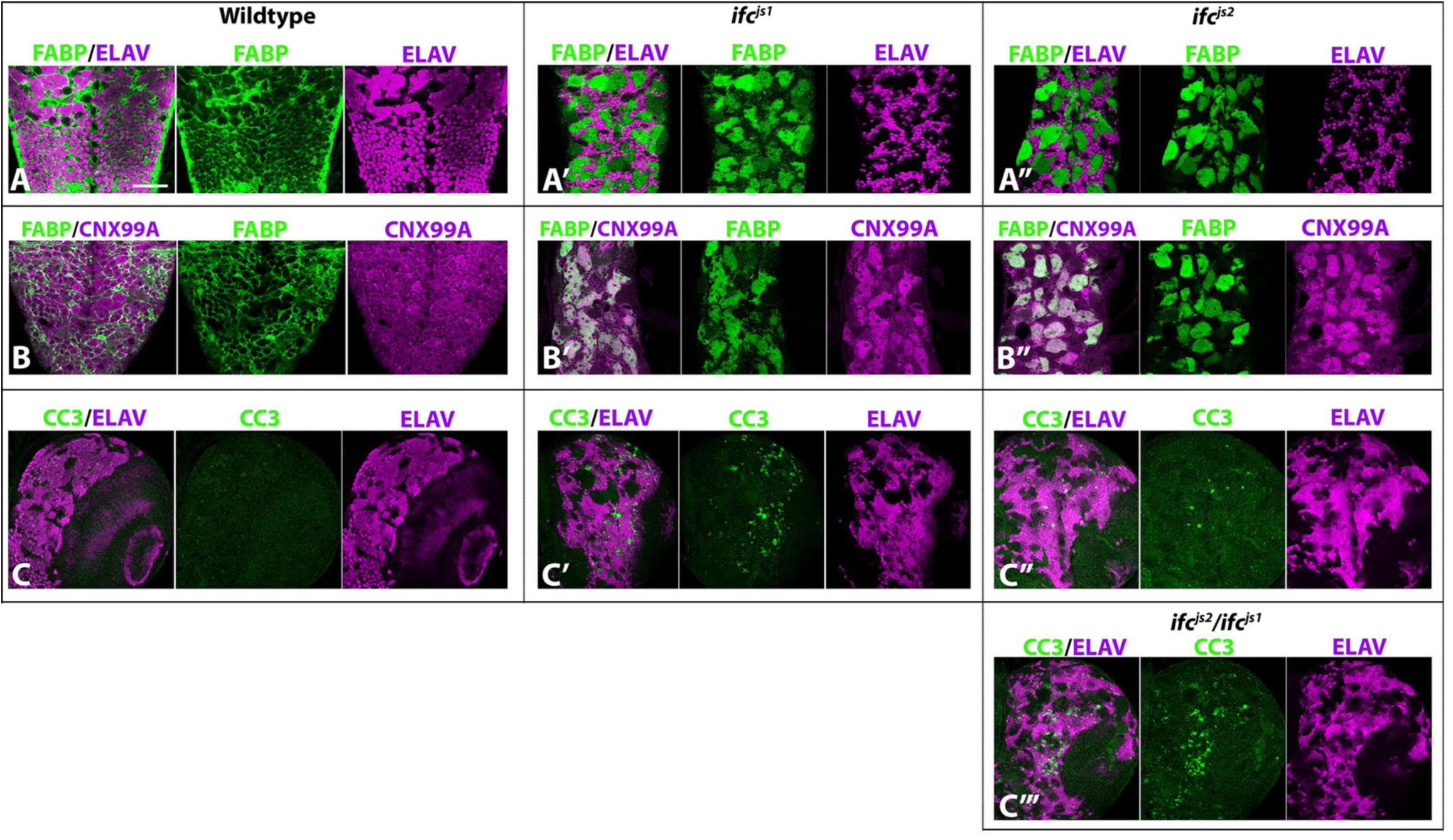
*ifc^js1^* and *ifc^js2^* alleles drive cortex glia swelling, ER expansion in cortex glia, and neuronal cell death. A-C) Abdominal segments of dissected ventral nerve cords from late-third instar larvae of the indicated genotype labeled for FABP to label cortex glia and ELAV to label neurons (A-A’’) or FABP and the ER-marker Calnexin-99A (CNX99A; B-B’’). As observed in *ifc^js3^*/*ifc-KO* larvae, *ifc^js1^* and *ifc^js2^* larvae exhibited swollen cortex glia (A-B) and an expanded ER phenotype in cortex glia as indicated by elevated CNX99A expression (B). (C-C’’’) *ifc^js1^* and *ifc^js1^*/*ifc^js2^*larvae also exhibit enhanced presence of the neuronal cell death marker cleaved Caspase 3 (CC3). Note that the chromosome that harbors the *ifc^js2^* chromosome also harbors a mutation in the pro-apoptotic gene, *Dark*. Therefore, to track the impact of *ifc^js2^*on neuronal cell death, we assessed neuronal cell death in larvae trans-heterozygous for *ifc^js1^* and *ifc^js2^* (Civ). Scale bar is 50 microns.

**Figure S8).**
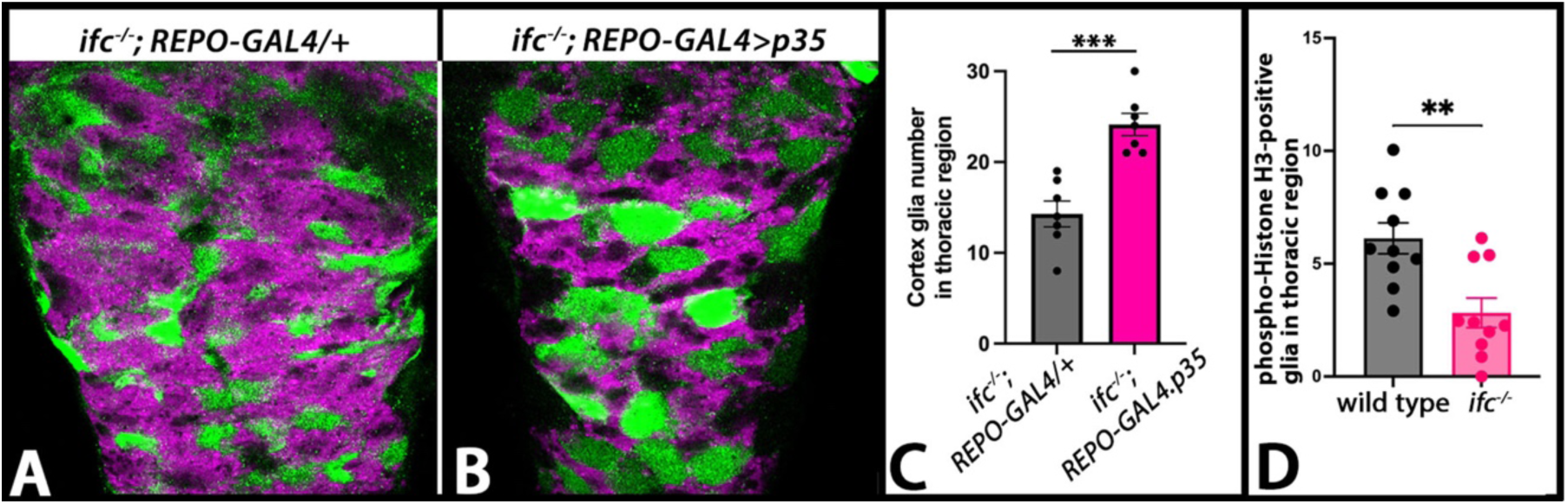
Reduction of cortex glia number in *ifc^-/-^* larvae results from increased apoptosis and reduced cell proliferation. A-B) Thoracic regions of the CNS of larvae of the indicated genotype labeled for ELAV (magenta) to label neurons and FABP (green) to label cortex glia. C) Number of cortex glia in the thoracic region of larvae of the indicated genotype; *** denotes p value less than 3 x 10^-4^. D) Number of phospho-Histone H3 positive glia in the thoracic region of larvae of the indicated genotype; ** denotes p value of less than 0.01.

**Figure S9.**
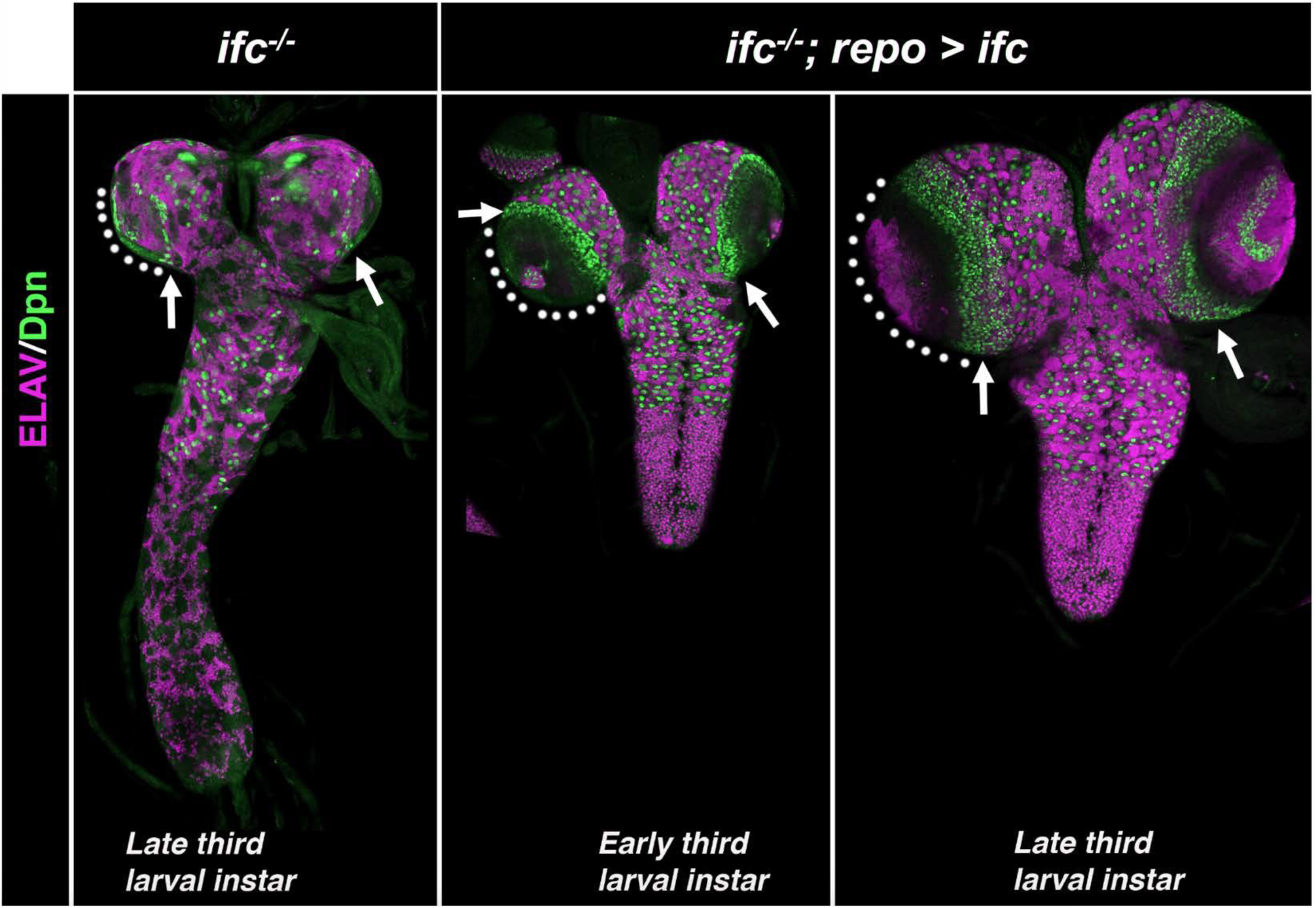
Glial-specific expression of *ifc* completely rescues the CNS phenotype of *ifc* mutant larvae. Ventral views of the CNS from *ifc* mutant larvae (left) and *ifc* mutant larvae in which a wild-type *ifc* transgene is driven specifically in glia under the control of *repo-GAL4* (middle and right panels). Each CNS is labeled for neuroblasts in green (DPN) and neurons in red (ELAV). Note the strong rescue of the CNS elongation and reduction of optic lobe size (dotted circles) and neuroblast number (arrows) observed in *ifc* mutant larvae relative to those that express *ifc* in glia. Anterior is up; scale bar is 50 microns.

**Figure S10.**
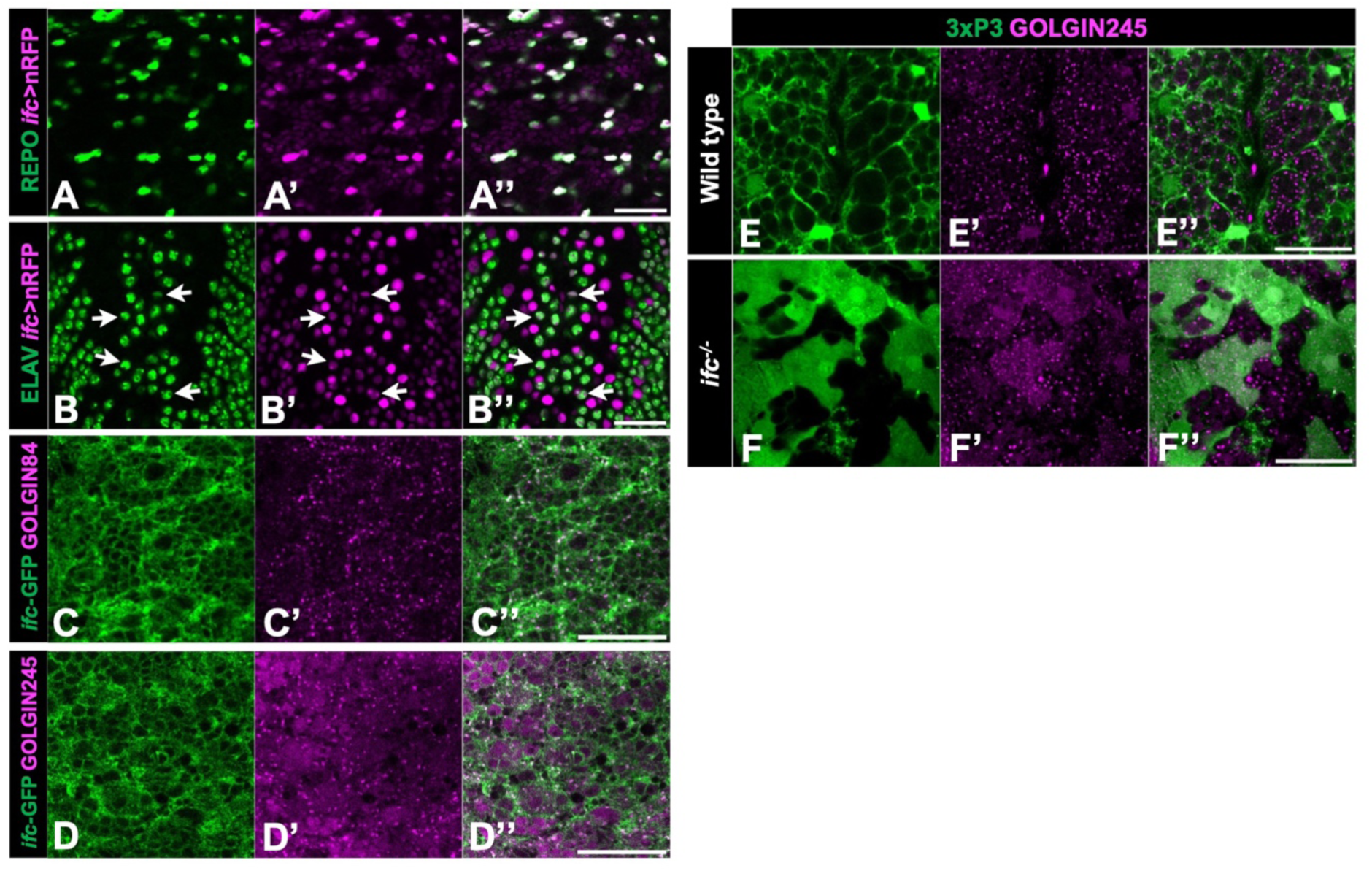
*ifc* is predominately expressed in glia and Ifc protein mildly localizes to the Golgi where mild expansion appears to occur in the larval CNS. A-B) Ventral (A-A’’) and dorsal (B-B’’) views of the CNS of late-third instar wild-type larvae labeled for *ifc-GAL4>nRFP* (magenta; A’-B’), REPO to mark glia (green; A), and ELAV to mark neurons (green; B). C-D) High magnification ventral views of thoracic segments in the CNS of wild-type late third instar larvae labeled for GFP (green; C and D), GOLGIN84 (magenta; C’), GOLGIN245 (magenta; D’). E-F) Late third instar larvae labeled for 3xP3-RFP (green; E and F), GOLGIN245 (magenta; E’ and F’). Anterior is up; scale bar is 30μm for all panels.

**Figure S11.**
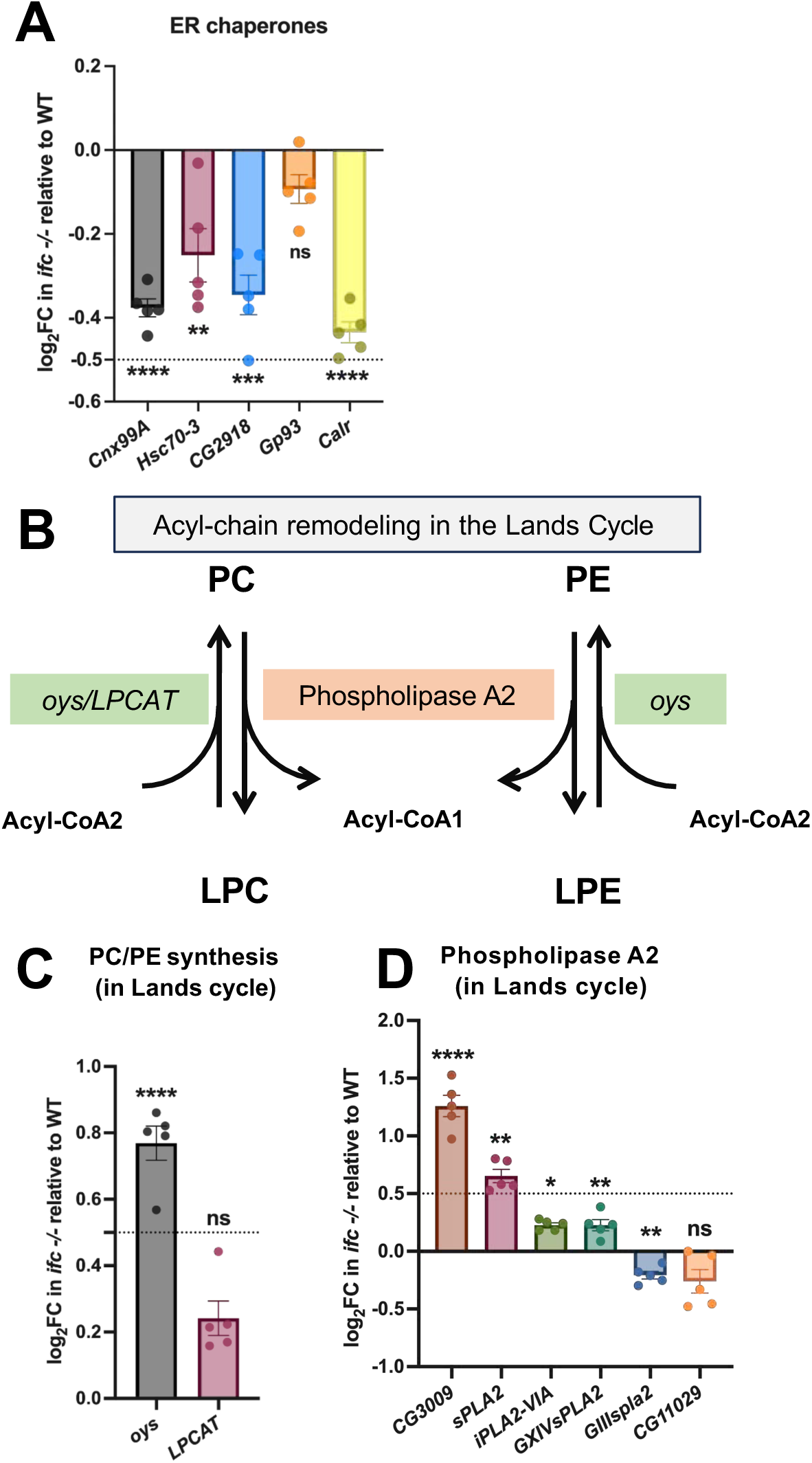
ER chaperones are mostly downregulated, and genes involved the Lands cycle are mostly upregulated in the CNS of *ifc^-/-^*mutant larvae. A) Four of the five ER chaperones in *Drosophila* (Schroder & Kaufman, 2006; Ryoo et al., 2007) are significantly downregulated in transcription, suggesting an atypical unfolded protein response (UPR) following the transcriptional upregulation of *Xbp-1s* in the *ifc*-deprived CNS of *ifc* mutant larvae. B) Schematic of the Lands cycle that functions in the acyl chain remodeling of membrane phospholipids. C-D) Most genes participating in the assembly (C) and removal (D) of acyl chains in the Lands cycle are significantly upregulated.

**Figure S12.**
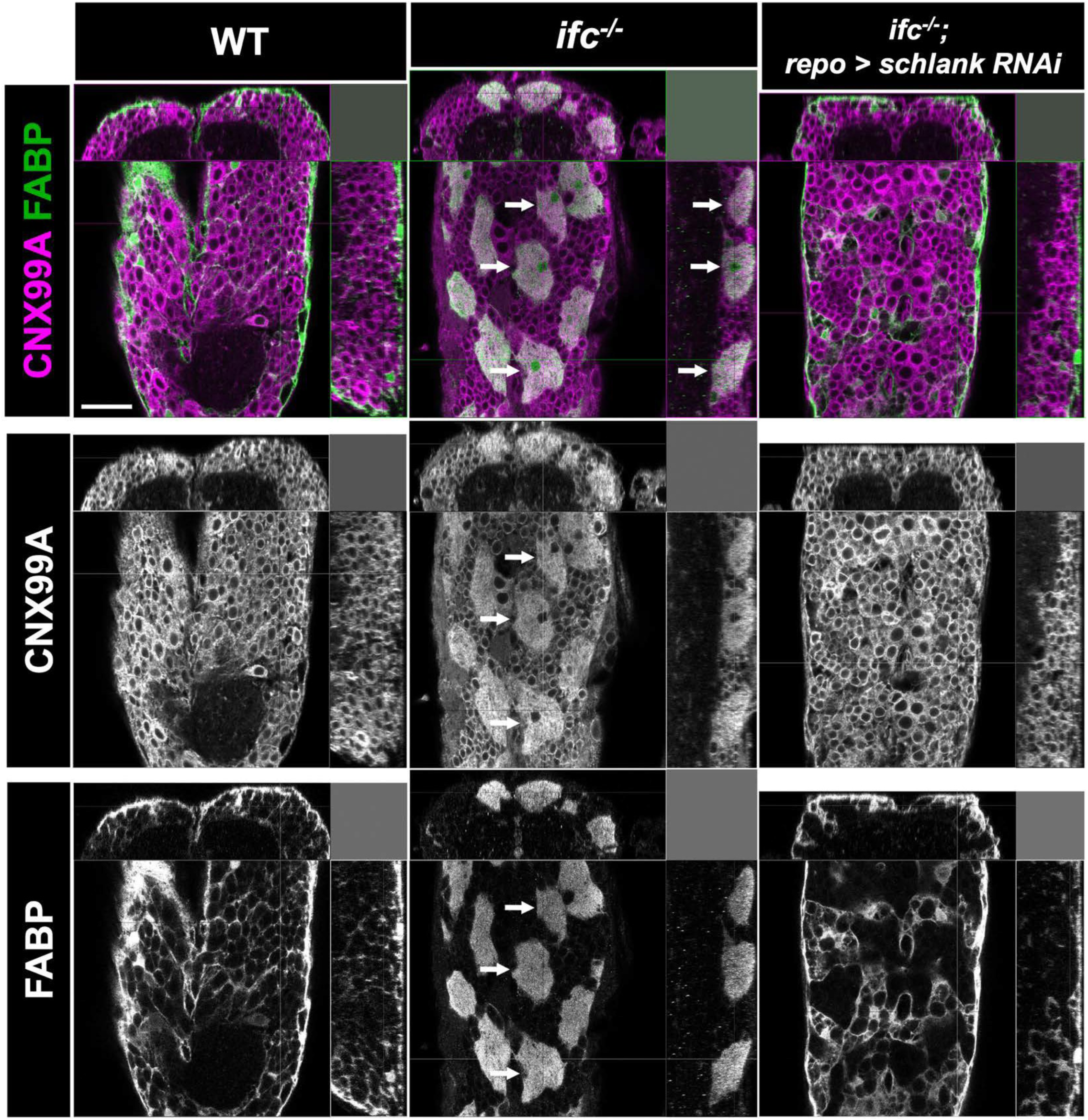
Glial-specific inhibition of *schlank* suppresses the *ifc* swollen cortex glia phenotype. High magnification X-Y, X-Z, and Y-Z views of the abdominal region of the ventral nerve cord of late third instar of the indicated genotype. FABP staining (green) labels cortex glia; CNX99A staining (red) labels ER. The swollen cortex glia in *ifc* mutant larvae (arrows) are not apparent in either wild-type larvae or *ifc* mutant larvae in which *schlank* function was inhibited specifically in glia (*ifc^-/-^; repo-GAL4/UAS-schlank^RNAi^*). Anterior is up; scale bar is 20μm.

**Figure S13.**
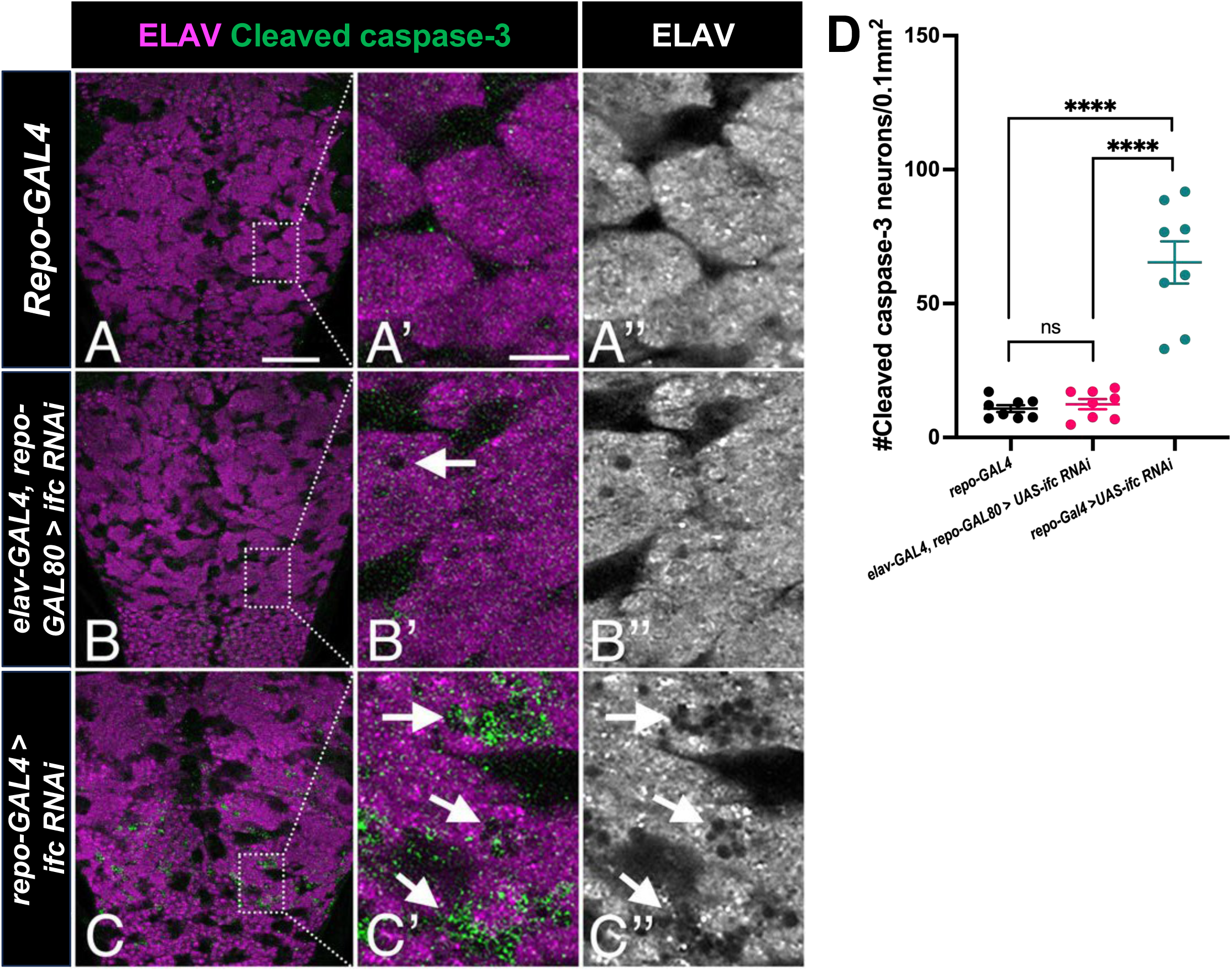
Glial-specific, but not neuronal-specific, knockdown of *ifc* drives neuronal cell death. Low (A-C) and high (A’-C’’) magnification views of the ventral nerve cord of late-third instar larvae of the indicated genotype labeled for ELAV (magenta or greyscale) and Cleaved Caspase-3 (green). Arrows point to regions marked by Cleaved Caspase-3 staining and/or apparent neuronal cell death identified by circular perforations in the neuronal cell layer. Please note that the *repoGAL80* transgene is present in *elav>ifc RNAi* larvae to block GAL4 function in glia (panel B-B’’). Anterior is up in all panels; scale bar is 50μm for panels A-C and 10μm for panels A’-C’’. Full genotypes of the larvae used in this experiment are as follows – A: *UAS-ifc-RNAi/+*; B: *elavGAL4/+; repoGAL80/+; UAS-ifc-RNAi/+*; C: *repoGAL4/UAS-ifc-RNAi*. D) Quantification of the number of Cleaved Caspase-3 positive neurons in the thorax region of the ventral nerve cord in control, neuronal-specific knockdown of *ifc*, and glial-specific knockdown of *ifc* L3 larvae.

**Figure S14.**
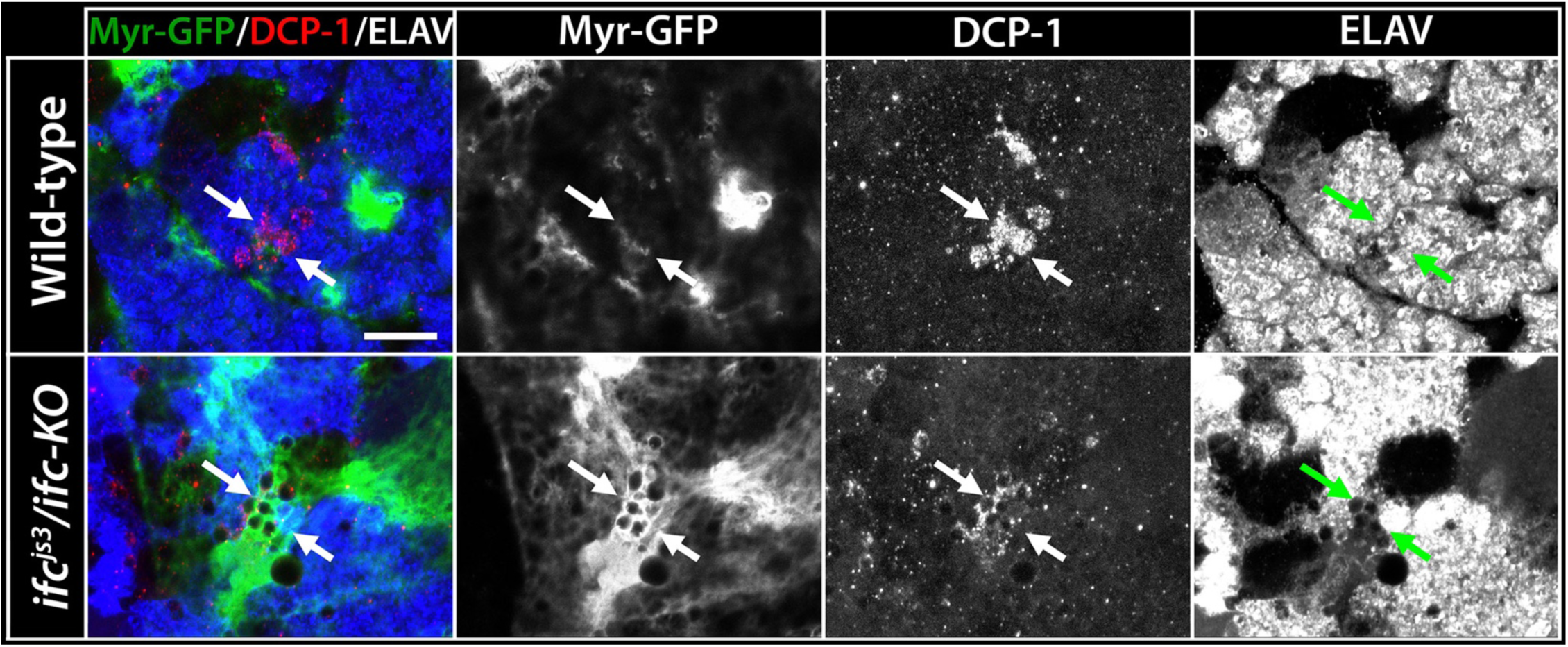
Subperineurial glial cell membranes encircle dying neurons in *ifc* mutant larvae. High magnification ventral views of the brain of wild-type (top) and *ifc* mutant (bottom) larvae labeled for Myr-GFP to mark the membranes of subperineurial glia (green, left panels), Death caspase protein-1 (DCP-1) to label dying neurons (red, left panels), and ELAV to mark neurons and the neuronal cell layer (blue, left panels). Arrows identify DCP-1 positive dying neurons, which are encircled by GFP-positive glial cell membranes in *ifc* mutant, but not wild-type, backgrounds. Anterior is up; scale bar is 10 microns. DCP-1 antibody obtained from Cell Signaling Technologies (Ab #9578)

**Table 1.** Genotype of flies in figures 2 & 3.

| Figure | Glial subtype | Group | Genotype |
| --- | --- | --- | --- |
| Figure 2 (a) | Cortex glia | Wild type | *GMR 54HO2-GAL4/10XUAS-IVS-myr-GFP* |
| Figure 2 (a') | Cortex glia | *ifc*<sup>-/-</sup> mutant | *ifc-KO/ifc<sup>js3</sup>; GMR 54HO2-GAL4/10XUAS-IVS-myr-GFP* |
| Figure 2 (b) | Ensheathing glia | Wild type | *GMR 56F03-GAL4/10XUAS-IVS-myr-GFP* |
| Figure 2 (b') | Ensheathing glia | *ifc*<sup>-/-</sup> mutant | *ifc-KO/ifc<sup>js3</sup>; GMR 56F03-GAL4/10XUAS-IVS-myr-GFP* |
| Figure 2 (c) | Astrocyte-like glia | Wild type | *GMR86EO1-GAL4/10XUAS-IVS-myr-GFP* |
| Figure 2 (c') | Astrocyte-like glia | *ifc*<sup>-/-</sup> mutant | *ifc-KO/ifc<sup>js3</sup>; GMR 56F03-GAL4/10XUAS-IVS-myr-GFP* |
| Figure 2 (d) | Subperineurial glia | Wild type | *GMR 54C07-GAL4/10XUAS-IVS-myr-GFP* |
| Figure 2 (d') | Subperineurial glia | *ifc*<sup>-/-</sup> mutant | *ifc-KO/ifc<sup>js3</sup>; GMR 54C07-GAL4/10XUAS-IVS-myr-GFP* |
| Figure 2 (e) | Perineurial glia | Wild type | *GMR 85G01-GAL4/10XUAS-IVS-myr-GFP* |
| Figure 2 (e') | Perineurial glia | *ifc*<sup>-/-</sup> mutant | *ifc-KO/ifc<sup>js3</sup>; GMR 85G01-GAL4/10XUAS-IVS-myr-GFP* |
| Figure 3 (a) | Cortex glia | Wild type | *yw* |
| Figure 3 (b) | Cortex glia | *ifc*<sup>-/-</sup> mutant | *ifc<sup>js3</sup>/ifc-KO; repoGAL4/UAS-NLS-GFP* |
| Figure 3 (c) | Cortex glia | Pan-neuronal *ifc* RNAi | *elavGAL4/+; repoGAL80/+; UAS-ifc[RNAi]/+* |
| Figure 3 (d) | Cortex glia | Pan-glial *ifc* RNAi | *repoGAL4/UAS-ifc[RNAi]* |
| Figure 3 (e) | Cortex glia | UAS-*ifc* control in *ifc*<sup>-/-</sup> mutant | *ifc<sup>js3</sup>/ifc-KO; UAS-ifc* |
| Figure 3 (f) | Cortex glia | Pan-neuronal *ifc* rescue in *ifc*<sup>-/-</sup> mutant | *elavGAL4/+; ifc<sup>js3</sup>/ifc-KO repoGAL80; UAS-ifc/+* |
| Figure 3 (g) | Cortex glia | Pan-glial *ifc* rescue in *ifc*<sup>-/-</sup> mutant | *ifc<sup>js3</sup>/ifc-KO; repoGAL4/UAS-ifc* |
| Figure 3 (h) | Cortex glia | Pan-neuronal *DEGS1* rescue in *ifc*<sup>-/-</sup> mutant | *elavGAL4/+; ifc<sup>js3</sup>/ifc-KO repoGAL80; UAS-DEGS1/+* |
| Figure 3 (i) | Cortex glia | Pan-glial *DEGS1* rescue in *ifc*<sup>-/-</sup> mutant | *ifc<sup>js3</sup>/ifc-KO; repoGAL4/UAS-DEGS1* |

